# Spatial geometry of the Hypoxic niche predicts radiotherapy failure in colorectal cancer: a spatially explicit agent-based model

**DOI:** 10.1101/2025.04.19.649627

**Authors:** Sa Dharmasastha Karthikeya, Prathiba Jonnala

## Abstract

Radiotherapy failure in locally advanced colorectal cancer is mechanistically linked to tumour microenvironmental heterogeneity, yet the spatial determinants of immune clearance following fractionated irradiation remain poorly understood. Here we present a spatially explicit hybrid agent-based model of colorectal cancer response to short-course radiotherapy in which cellular fate—proliferation, dormancy, or necrosis—is governed by local oxygen tension and ATP availability computed via Michaelis–Menten kinetics. Immune killing probability and proliferation rate are implemented as continuous functions of local oxygen, replacing hard thresholds with mechanistically grounded gradients. Simulations of the standard 5 *×* 5 Gy short-course radiotherapy protocol revealed a bimodal tumour control distribution consistent with clinical observations of cure-or-escape outcomes. The central finding is that spatial position within the oxygen landscape, rather than surviving cell count alone, predicts immune clearance success: runs in which the tumour was eliminated had surviving cells in well-oxygenated regions near vasculature (mean O_2_ = 0.967), while escaped tumours retained cells concentrated in the hypoxic core (mean O_2_ = 0.196; Mann–Whitney *p* = 2.1 *×* 10*^−^*^9^). Fractionation schedule analysis across equal total-dose protocols demonstrated that inter-fraction interval critically modulates repopulation dynamics and tumour control probability, with compact schedules outperforming widelyspaced fractionation despite identical cumulative dose. Transcriptomic validation using the Buffa hypoxia signature across 284 Stage II/III CRC patients from TCGA confirmed that significant tumour hypoxia is a clinically prevalent feature of the target disease population. These results suggest that the geometry of the hypoxic niche is a spatial determinant of radiotherapy outcome in colorectal cancer, with implications for treatment scheduling and patient stratification by hypoxia burden.

**Highlights:**

- Spatial position of surviving cells within the oxygen landscape, not surviving cell count, predicts immune clearance success following fractionated radiotherapy.
- A hybrid agent-based model of colorectal cancer demonstrates complete outcome separation by mean survivor oxygen concentration across 50 independent realisations.
- Inter-fraction interval critically modulates tumour control probability under equal total-dose conditions; hypofractionation outperforms daily fractionation through repopulation suppression.
- Transcriptomic analysis of 284 Stage II/III CRC patients confirms clinically prevalent intratumoral hypoxia, grounding model oxygen assumptions in real patient biology.
- The hypoxic niche functions as a microenvironmental refuge through coincident suppression of radiation efficacy and immune killing probability by the same oxygen-dependent mechanism.

## 1. Introduction

Colorectal cancer (CRC) remains the third most commonly diagnosed malignancy and the second leading cause of cancer-related mortality worldwide, with locally advanced rectal cancer representing a clinically challenging subgroup in which achieving locoregional control is critical to long-term survival [1, 2]. For Stage II and III disease, neoadjuvant radiotherapy—delivered prior to surgical resection—is a standard component of multimodal treatment, with short-course radiotherapy (SCRT) delivering 25 Gy in five daily fractions emerging as an evidence-based alternative to prolonged chemoradiotherapy schedules [3, 4, 5]. Beyond its direct cytotoxic effects, radiation is now understood to engage the immune system in ways that extend beyond the irradiated field. Ionising radiation induces immunogenic cell death, promoting the release of damage-associated molecular patterns and tumour-associated antigens that recruit and activate cytotoxic T lymphocytes (CTLs) [6, 7]. This radiation–immune synergy has motivated growing interest in combined RT–immunotherapy strategies for CRC, particularly in mismatch repair-deficient tumours [8, 9]. However, the conditions under which this synergy succeeds or fails in the spatial tumour microenvironment remain incompletely understood.

A well-established but underappreciated obstacle to both radiotherapy efficacy and immune-mediated tumour clearance is intratumoral hypoxia. Solid tumours frequently develop regions of severe oxygen deficiency as proliferating cells outpace the diffusion capacity of an irregular and inadequate vasculature [10, 11, 12]. In these hypoxic zones, radiation is substantially less effective: the oxygen enhancement ratio (OER) quantifies this dependence, with anoxic cells requiring approximately 2.5-fold higher doses to achieve equivalent cytotoxicity compared to well-oxygenated cells [13, 14]. Simultaneously, hypoxia impairs the effector function of tumour-infiltrating CTLs through multiple mechanisms including HIF-1*α*-mediated upregulation of immune checkpoint ligands, adenosine accumulation, and direct suppression of granzyme-B activity [15, 16]. The consequence is a spatially defined region within the tumour—the hypoxic niche—where cells are simultaneously protected from radiation and shielded from immune killing. Metabolically, cells within this niche sustain survival through anaerobic glycolysis despite severely reduced oxidative phosphorylation, a phenotypic state characterised by the Warburg effect [17, 18, 19]. These cells are not simply radioresistant; they occupy a microenvironmental refuge that renders standard treatment combinations insufficient for their elimination.

Despite this mechanistic understanding, existing mathematical models of CRC response to radiotherapy have largely treated the tumour as a spatially uniform population. Ordinary differential equation frameworks and population-averaged linear-quadratic models capture bulk tumour kinetics but cannot represent the spatial oxygen gradients, local metabolic states, or positional relationships between surviving cells and immune effectors that determine whether post-irradiation clearance succeeds [20, 21, 22]. The spatial question—specifically, whether the location of radiotherapy survivors within the oxygen landscape predicts immune clearance outcome—has not been systematically addressed in a mechanistic computational framework for CRC. This gap is clinically significant. If treatment failure is determined not by how many cells survive radiation but by *where* they survive, then patient stratification by tumour hypoxia burden and treatment schedule optimisation around immune–RT timing become mechanistically motivated rather than empirically derived strategies.

Here we present a spatially explicit hybrid agent-based model of colorectal cancer responding to fractionated short-course radiotherapy with concurrent immune feedback. Each cell agent occupies a discrete lattice site and its behaviour—proliferation, metabolic dormancy, or necrosis—is governed entirely by local oxygen tension and ATP availability, computed through Michaelis–Menten kinetics from a solved oxygen diffusion field. Radiotherapy is applied as an oxygen enhancement ratio-corrected linear-quadratic survival probability per cell, and CTL killing probability is implemented as a continuous function of local oxygen rather than a population-averaged thresh-old. We use this model to test the hypothesis that spatial geometry of the hypoxic niche, rather than aggregate surviving cell burden, is the primary determinant of whether immune clearance succeeds following fractionated radiotherapy in colorectal cancer.

## 2. Methods

### 2.1. Model Overview and Lattice Geometry

We developed a two-dimensional hybrid agent-based model in which in-dividual tumour cells occupy discrete sites on a 50 *×* 50 lattice representing a 1 mm^2^ cross-section of colorectal tumour tissue, with each lattice site corresponding to a 20 *µ*m *×* 20 *µ*m domain consistent with the diameter of a single epithelial cell. Each site is classified at every timestep as one of fivestates: empty, sensitive (quiescent), proliferative, resistant (mutant), or decaying. The simulation advances in hourly timesteps over a total duration of 240 hours, spanning the full 5-fraction radiotherapy course and a 144-hour post-treatment observation window. All cellular behaviours—proliferation rate, radiosensitivity, and susceptibility to immune killing—are determined exclusively by local microenvironmental conditions computed at each site, with no global or population-averaged parameters governing individual cell fate.

**Table 1:**
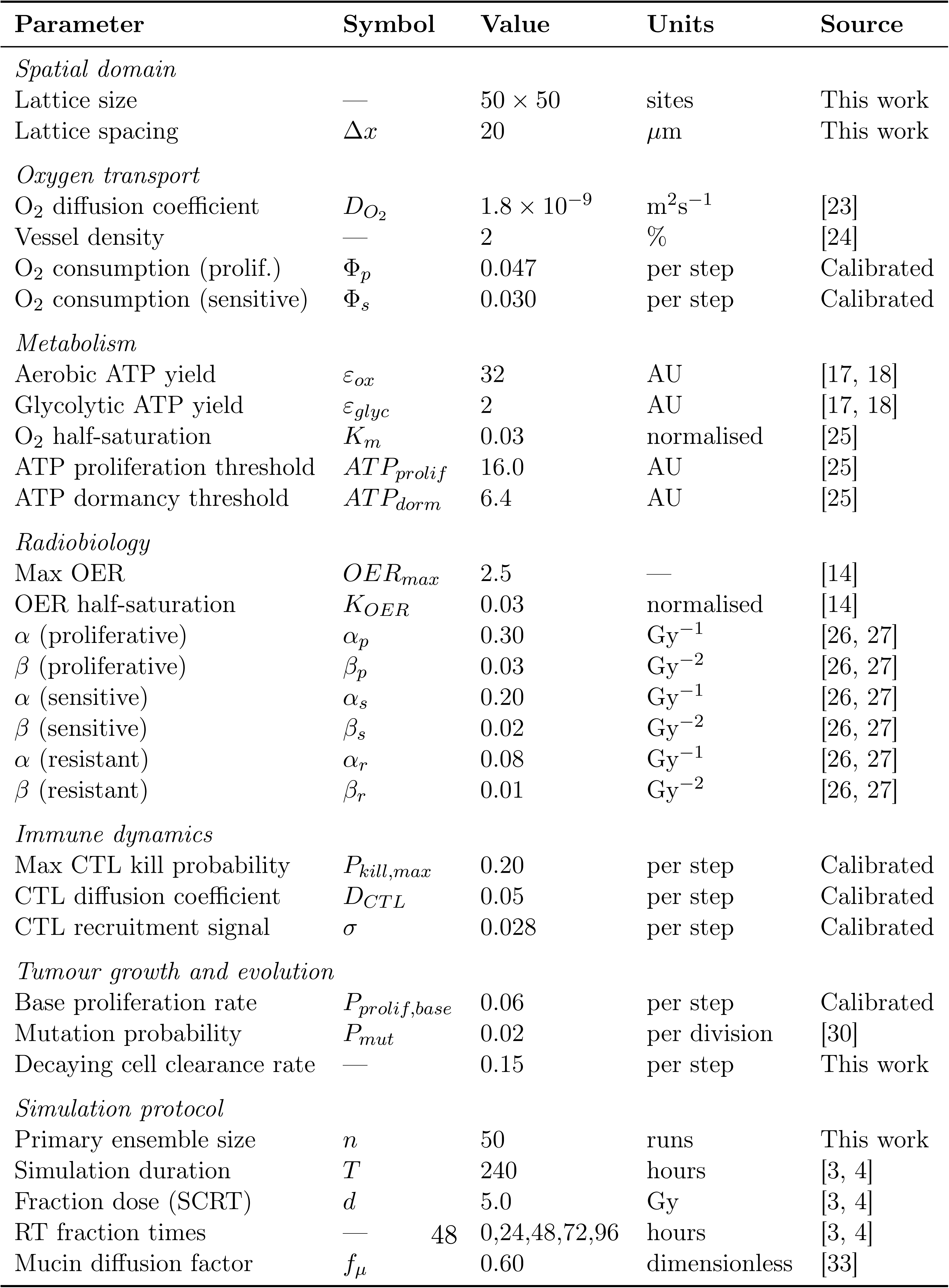
Model parameters, symbols, values, units, and sources. AU = arbitrary units (normalised ATP equivalents); per step = per hourly simulation timestep.

| **Parameter** | **Symbol** | **Value** | **Units** | **Source** |
|---|---|---|---|---|
| *Spatial domain* |  |  |  |  |
| Lattice size | — | 50 × 50 | sites | This work |
| Lattice spacing | Δx | 20 | μm | This work |
| *Oxygen transport* |  |  |  |  |
| O<sub>2</sub> diffusion coefficient | D<sub>O<sub>2</sub></sub> | 1.8 × 10<sup>−9</sup> | m<sup>2</sup>s<sup>−1</sup> | [23] |
| Vessel density | — | 2 | % | [24] |
| O<sub>2</sub> consumption (prolif.) | Φ<sub>p</sub> | 0.047 | per step | Calibrated |
| O<sub>2</sub> consumption (sensitive) | Φ<sub>s</sub> | 0.030 | per step | Calibrated |
| *Metabolism* |  |  |  |  |
| Aerobic ATP yield | ε<sub>ox</sub> | 32 | AU | [17, 18] |
| Glycolytic ATP yield | ε<sub>glyc</sub> | 2 | AU | [17, 18] |
| O<sub>2</sub> half-saturation | K<sub>m</sub> | 0.03 | normalised | [25] |
| ATP proliferation threshold | ATP<sub>prolif</sub> | 16.0 | AU | [25] |
| ATP dormancy threshold | ATP<sub>dorm</sub> | 6.4 | AU | [25] |
| *Radiobiology* |  |  |  |  |
| Max OER | OER<sub>max</sub> | 2.5 | — | [14] |
| OER half-saturation | K<sub>OER</sub> | 0.03 | normalised | [14] |
| α (proliferative) | α<sub>p</sub> | 0.30 | Gy<sup>−1</sup> | [26, 27] |
| β (proliferative) | β<sub>p</sub> | 0.03 | Gy<sup>−2</sup> | [26, 27] |
| α (sensitive) | α<sub>s</sub> | 0.20 | Gy<sup>−1</sup> | [26, 27] |
| β (sensitive) | β<sub>s</sub> | 0.02 | Gy<sup>−2</sup> | [26, 27] |
| α (resistant) | α<sub>r</sub> | 0.08 | Gy<sup>−1</sup> | [26, 27] |
| β (resistant) | β<sub>r</sub> | 0.01 | Gy<sup>−2</sup> | [26, 27] |
| *Immune dynamics* |  |  |  |  |
| Max CTL kill probability | P<sub>kill,max</sub> | 0.20 | per step | Calibrated |
| CTL diffusion coefficient | D<sub>CTL</sub> | 0.05 | per step | Calibrated |
| CTL recruitment signal | σ | 0.028 | per step | Calibrated |
| *Tumour growth and evolution* |  |  |  |  |
| Base proliferation rate | P<sub>prolif,base</sub> | 0.06 | per step | Calibrated |
| Mutation probability | P<sub>mut</sub> | 0.02 | per division | [30] |
| Decaying cell clearance rate | — | 0.15 | per step | This work |
| *Simulation protocol* |  |  |  |  |
| Primary ensemble size | n | 50 | runs | This work |
| Simulation duration | T | 240 | hours | [3, 4] |
| Fraction dose (SCRT) | d | 5.0 | Gy | [3, 4] |
| RT fraction times | — 48 | 0,24,48,72,96 | hours | [3, 4] |
| Mucin diffusion factor | f<sub>μ</sub> | 0.60 | dimensionless | [33] |

### 2.2. Oxygen Diffusion and Vascular Architecture

Oxygen transport was modelled using explicit finite-difference solution of the two-dimensional diffusion equation with cellular consumption:

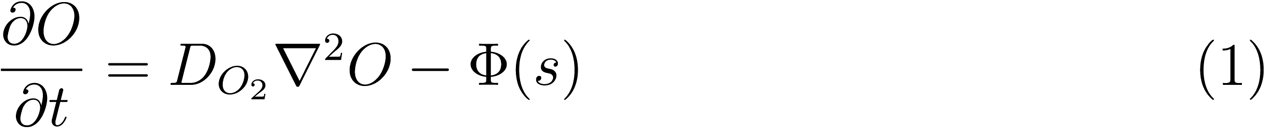

where *O* denotes normalised oxygen concentration (dimensionless, 0–1), *D_O_*_2_ = 1.8 *×* 10*^−^*^9^ m^2^s*^−^*^1^ is the oxygen diffusion coefficient in tumour tissue [23], and Φ(*s*) is the oxygen consumption rate at site *s*, set to 0.047 per timestep for proliferative cells and 0.030 per timestep for sensitive and resistant cells. Numerical stability was ensured by enforcing the von Neumann stability condition:

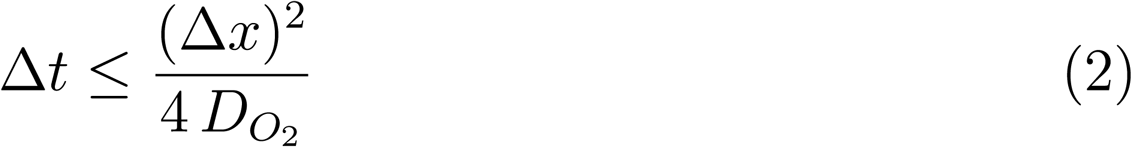

with Δ*x* = 20 *µ*m giving a stable diffusion substep of Δ*t*_diff_. Five diffusion substeps were computed per hourly biological timestep to approximate quasisteady oxygen equilibration. Vascular supply was represented by a fixed set of source sites, comprising 2% of lattice positions, assigned at simulation initialisation using a fixed random seed and held constant across all realisations. At each source site, oxygen concentration was reset to 1.0 (normoxia) after every diffusion substep, representing continuous supply from patent microvessels. This vessel density is consistent with reported microvessel density in Stage II/III CRC tissue [10, 24].

### 2.3. Cellular Metabolism: ATP and the Hypoxic Niche

Cellular energy availability at each lattice site was computed from local oxygen tension using a Michaelis–Menten formulation representing the competitive balance between oxidative phosphorylation and anaerobic glycolysis:

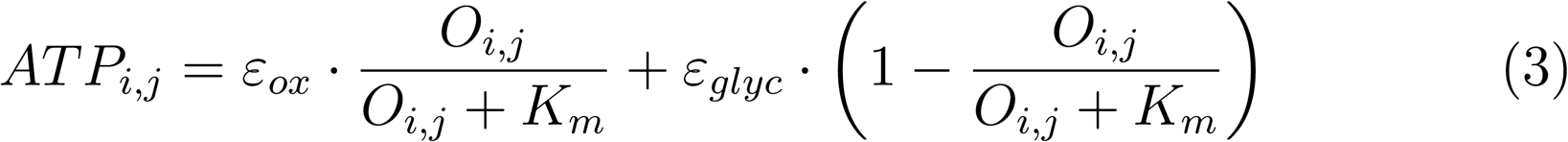

where *ε_ox_* = 32 ATP equivalents represents maximum aerobic yield, *ε_glyc_* = 2 ATP equivalents represents glycolytic yield consistent with the Warburg phenotype [17, 18], and *K_m_* = 0.03 (normalised units, corresponding to approximately 1.2 mmHg) is the oxygen half-saturation constant. This formulation produces a continuous metabolic transition from oxidative to glycolytic metabolism as oxygen falls below *K_m_*, without imposing a discrete threshold. Cellular metabolic state was classified according to local ATP level at each timestep. Sites with *ATP ≥* 16 (that is, *ATP ≥* 0.5 *· ε_ox_*) were designated proliferative, with sufficient energy for active division. Sites with 6.4 *≤ ATP <* 16 (corresponding to 0.2 *· ε_ox_≤ ATP <* 0.5 *· ε_ox_*) were designated metabolically dormant—viable but energetically unable to divide, constituting the hypoxic niche modelled in this work. Sites with *ATP <* 6.4 underwent necrotic removal from the lattice. These thresholds are consistent with published energy requirements for mammalian cell cycle progression [25, 18].

### 2.4. Radiotherapy Implementation

Radiotherapy fractions were applied at hours 0, 24, 48, 72, and 96, consistent with the standard 5 *×* 5 Gy short-course radiotherapy protocol for rectal cancer [3, 4]. At each fraction, the survival probability of every occupied lattice site was computed using the linear-quadratic model with oxygen enhancement ratio correction:

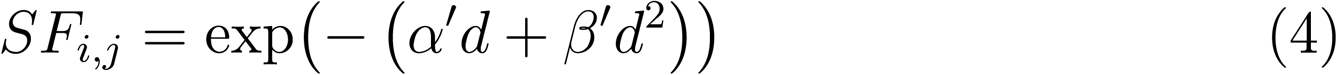

where *d* = 5 Gy is the fraction dose and the effective radiosensitivity parameters are:

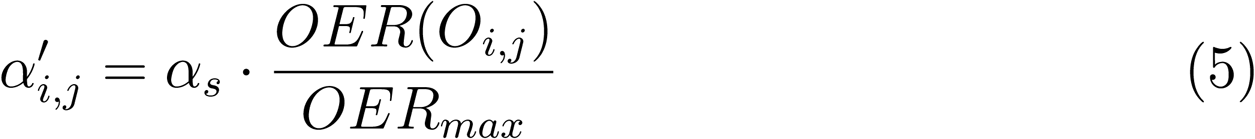

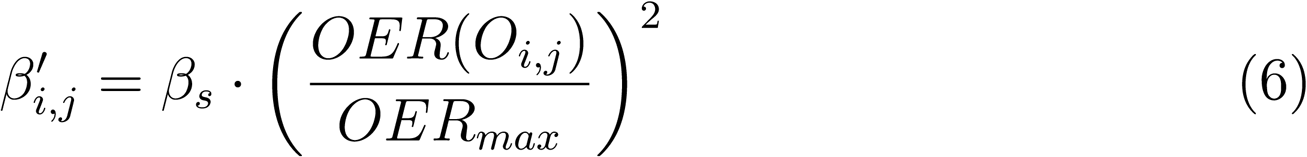

The oxygen enhancement ratio at each site was computed using the Alper– Howard–Flanders formulation [14]:

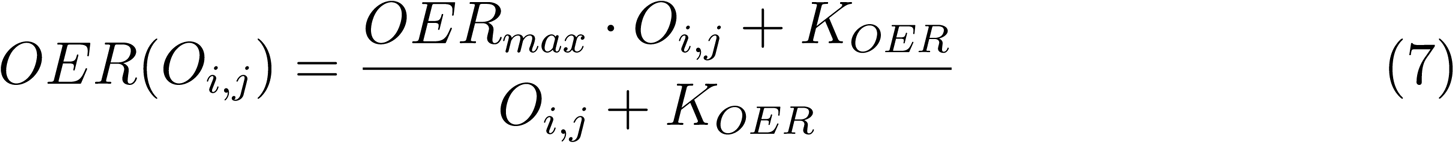

with *OER_max_*= 2.5 and *K_OER_* = 0.03. Radiosensitivity parameters were assigned by cell state: proliferative cells (*α* = 0.30 Gy*^−^*^1^, *β* = 0.03 Gy*^−^*^2^), sensitive cells (*α* = 0.20 Gy*^−^*^1^, *β* = 0.02 Gy*^−^*^2^), and resistant mutant cells (*α* = 0.08 Gy*^−^*^1^, *β* = 0.01 Gy*^−^*^2^), consistent with published *α/β* ratios for colorectal cancer [26, 27, 28, 13]. Cell fate following each fraction was determined by an independent Bernoulli trial: sites where rand() *> SF_i,j_*transitioned to decaying state. The calibration of the radiation physics implementation was confirmed by computing the population-averaged survival fraction following a single 5 Gy fraction (*SF*_5_ = 0.323), which reflects the spatially averaged response across the full oxygen distribution including the hypoxic core, and comparing this to the normoxic LQ prediction for sensitive cells (*SF*_5_ = 0.223); the difference is attributable to the radioresistant hypoxic subpopulation, consistent with published observations of hypoxia-driven radiosensitivity heterogeneity in solid tumours [13, 29].

### 2.5. Immune Dynamics

Cytotoxic T lymphocyte activity was represented by a continuous CTL density field *C_i,j_*(*t*) updated at each timestep according to:

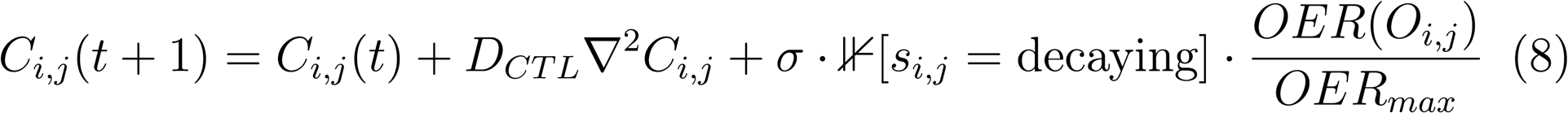

where *D_CT L_*= 0.05 is the CTL diffusion coefficient, *σ* = 0.028 is the recruitment signal strength per decaying cell (calibrated to produce a tumour control probability consistent with clinical SCRT outcomes), and the final term implements continuous hypoxic suppression of CTL recruitment— regions with low oxygen simultaneously generate weaker recruitment signals. CTL density was clipped to [0, 1] at each timestep.

The probability of CTL-mediated killing of an occupied site was implemented as a continuous function of local oxygen:

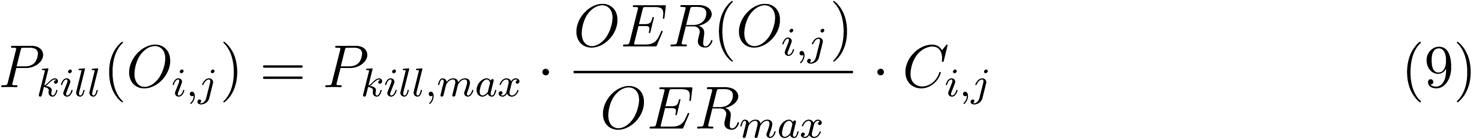

with *P_kill,max_* = 0.20. This formulation ensures that killing probability approaches zero under anoxia and reaches its maximum under normoxia, capturing the documented impairment of CTL effector function in hypoxic tumour microenvironments [15, 16]. This replaces the binary killing thresh-old used in earlier implementations [20], producing mechanistically grounded spatial immune suppression without discontinuities.

### 2.6. Cell Fate Rules

At each hourly timestep, following oxygen diffusion and immune updates, cell fate was determined by local ATP. Cells with *ATP <* 6.4 were removed from the lattice. Cells with 6.4 *≤ ATP <* 16 remained viable but did not divide. Cells with *ATP ≥* 16 entered the proliferative state, with division probability:

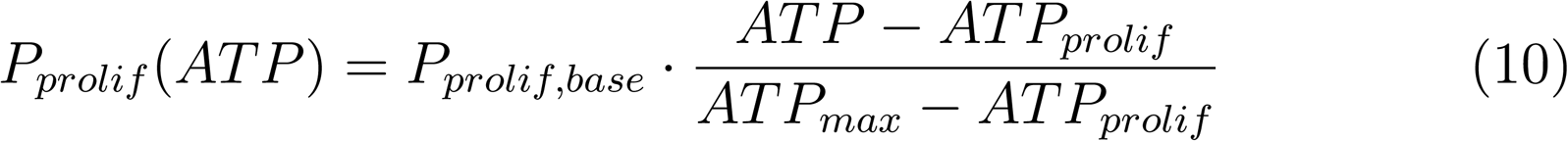

where *P_prolif,base_* = 0.06 and *ATP_prolif_* = 16. Division placed a daughter cell in a randomly selected empty Moore neighbour. With probability *P_mut_* = 0.02 per division, a daughter cell acquired the resistant phenotype with reduced radiosensitivity, following the stochastic resistance emergence framework of Goldie and Coldman [30]. Decaying cells were cleared from the lattice stochastically at a rate of 0.15 per timestep.

### 2.7. Simulation Protocols and Ensemble Design

The primary simulation protocol followed the standard 5 *×* 5 Gy SCRT schedule with concurrent immune response. All results presented in the primary analysis derive from an ensemble of 50 independent realisations, each initialised with a distinct random seed while maintaining identical model parameters and vascular geometry. Ensemble size was determined by bootstrap convergence analysis: TCP estimates stabilised at *n ≥* 25, with 90% bootstrap confidence interval width falling below 10% at *n* = 45 (Supplementary convergence analysis). A parallel ensemble of 50 realisations without immune response was run under identical conditions to isolate the contribution of immune dynamics to tumour control.

Protocol comparison was conducted across three equal total-dose schedules, each delivering 25 Gy: the standard SCRT protocol (5 *×* 5 Gy, fractions at 0, 24, 48, 72, 96 hours), a widely-spaced fractionation schedule (5 *×* 5 Gy, fractions at 0, 48, 96, 144, 192 hours), and a hypofractionation protocol (2 *×* 12.5 Gy, fractions at 0 and 48 hours). Protocol comparison ensembles used 30 realisations per arm, sufficient given demonstrated convergence by *n* = 25. All three protocols included immune response. Equal total dose was maintained across protocols to isolate fractionation schedule effects from dose effects.

Global sensitivity analysis was performed using Latin Hypercube Sampling across four parameters: radiosensitivity (*α*, range 0.10–0.40 Gy*^−^*^1^), base proliferation rate (*P_prolif,base_*, range 0.03–0.10), maximum CTL killing probability (*P_kill,max_*, range 0.08–0.35), and oxygen half-saturation constant (*K_m_*, range 0.015–0.060). Forty parameter sets were sampled, each evaluated with three independent realisations (120 total model evaluations), following established practice for agent-based model sensitivity analysis [31]. Pearson correlation coefficients between each parameter and final tumour burden were computed across the 40 sample means.

The justification for differing ensemble sizes across analyses (*n* = 50 primary, *n* = 40 GSA, *n* = 30 protocol comparison) is as follows: the primary ensemble maximises statistical power for the spatial clustering analysis, which is the central result; the GSA measures parameter–outcome correlations across 40 LHS samples with three replicates each (120 total evaluations), where the relevant unit is the LHS sample not the individual realisation; and protocol comparison uses *n* = 30 per arm, sufficient given convergence analysis and chosen for computational tractability across three parallel arms.

**Algorithm 1:**
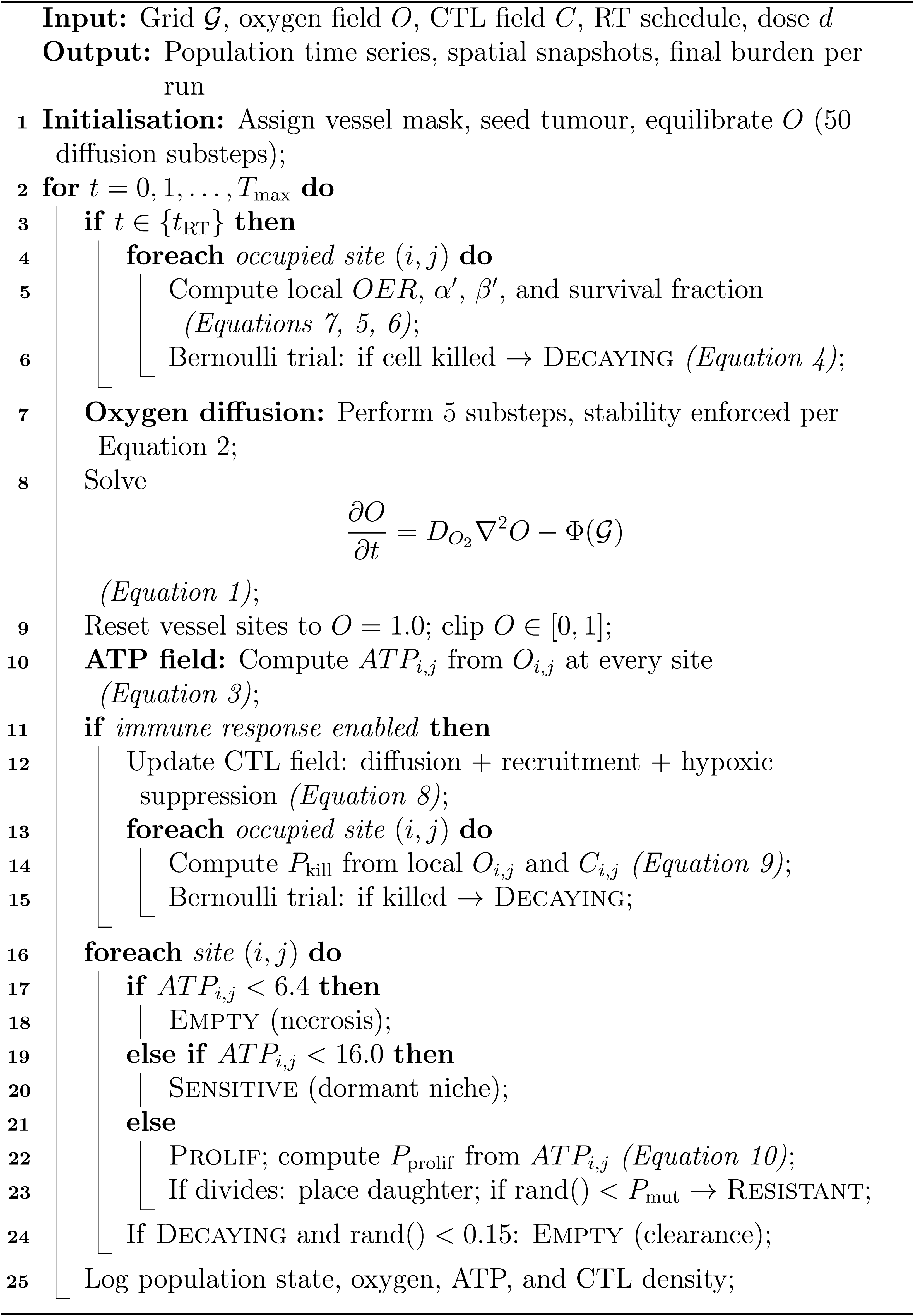
CRC Hybrid ABM — Main Simulation Loop.

|  |  |
|---|---|
|  | **Input:** Grid 𝒢, oxygen field O, CTL field C, RT schedule, dose d |
|  | **Output:** Population time series, spatial snapshots, final burden per run |
| 1 | **Initialisation:** Assign vessel mask, seed tumour, equilibrate O (50 diffusion substeps); |
| 2 | **for** t = 0, 1, . . . , T<sub>max</sub> **do** |
| 3 | **if** t ∈ {t<sub>RT</sub>} **then** |
| 4 | **foreach** *occupied site* (i, j) **do** |
| 5 | Compute local OER, α′, β′, and survival fraction *(Equations 7, 5, 6)*; |
| 6 | Bernoulli trial: if cell killed → Decaying *(Equation 4)*; |
| 7 | **Oxygen diffusion:** Perform 5 substeps, stability enforced per Equation 2; |
| 8 | Solve<br>$\frac{\partial O}{\partial t} = D_{O_2}\nabla^2 O - \Phi(\mathcal{G})$<br>*(Equation 1)*; |
| 9 | Reset vessel sites to O = 1.0; clip O ∈ [0, 1]; |
| 10 | **ATP field:** Compute ATP<sub>i,j</sub> from O<sub>i,j</sub> at every site *(Equation 3)*; |
| 11 | **if** *immune response enabled* **then** |
| 12 | Update CTL field: diffusion + recruitment + hypoxic suppression *(Equation 8)*; |
| 13 | **foreach** *occupied site* (i, j) **do** |
| 14 | Compute P<sub>kill</sub> from local O<sub>i,j</sub> and C<sub>i,j</sub> *(Equation 9)*; |
| 15 | Bernoulli trial: if killed → Decaying; |
| 16 | **foreach** *site* (i, j) **do** |
| 17 | **if** ATP<sub>i,j</sub> < 6.4 **then** |
| 18 | Empty (necrosis); |
| 19 | **else if** ATP<sub>i,j</sub> < 16.0 **then** |
| 20 | Sensitive (dormant niche); |
| 21 | **else** |
| 22 | Prolif; compute P<sub>prolif</sub> from ATP<sub>i,j</sub> *(Equation 10)*; |
| 23 | If divides: place daughter; if rand() < P<sub>mut</sub> → Resistant; |
| 24 | If Decaying and rand() < 0.15: Empty (clearance); |
| 25 | Log population state, oxygen, ATP, and CTL density; |

### 2.8. Transcriptomic Validation Cohort

To assess whether the oxygen microenvironment modelled in the ABM reflects clinically documented hypoxia in Stage II/III colorectal cancer, tumour hypoxia burden was quantified in 284 patients from the TCGA-COAD and TCGA-READ cohorts with available RNA sequencing data matched to clinical stage annotation (Stage II/III combined). Hypoxia scores were computed for each patient using single-sample gene set enrichment analysis applied to the Buffa 15-gene hypoxia signature [32]. Patients were classified as high-hypoxia if their ssGSEA score exceeded the 80th percentile threshold, consistent with the stratification approach defined in the original signature validation [32]. This analysis was performed in R using the GSVA package. The resulting hypoxia distribution provides biological justification for incorporating spatial oxygen heterogeneity as a mechanistic feature of the model rather than an ungrounded numerical assumption.

### 2.9. Statistical Analysis

Differences in final tumour burden between RT-only and RT-plus-immune ensembles were assessed using the two-sided Mann–Whitney *U* test, appropriate for non-normally distributed count data. Differences in mean oxygen concentration of surviving cells between cleared and escaped simulation runs were assessed by the same method. Tumour control probability was computed as the fraction of ensemble realisations in which final live cell count fell below five cells, with 95% confidence intervals derived using the Wilson score method. Pearson correlation coefficients were reported for global sensitivity analysis. All analyses were implemented in Python 3 using SciPy, NumPy, and statsmodels. Statistical significance was set at *α* = 0.05. Transcriptomic analysis was performed in the R computational environment using the GSVA package for ssGSEA scoring.

### 2.10. CRC-specific microenvironmental validation: mucin-mediated oxygen diffusion

Colorectal adenocarcinomas are characterised by mucin secretion — a thick glycoprotein layer that reduces effective oxygen diffusivity within the tumour mass relative to surrounding normal tissue [33]. To assess whether the central finding of this study is robust to this CRC-specific microenvironmental feature, we implemented a spatially varying oxygen diffusion coefficient in which the effective diffusivity within the tumour boundary was reduced to 60% of the free-tissue value:

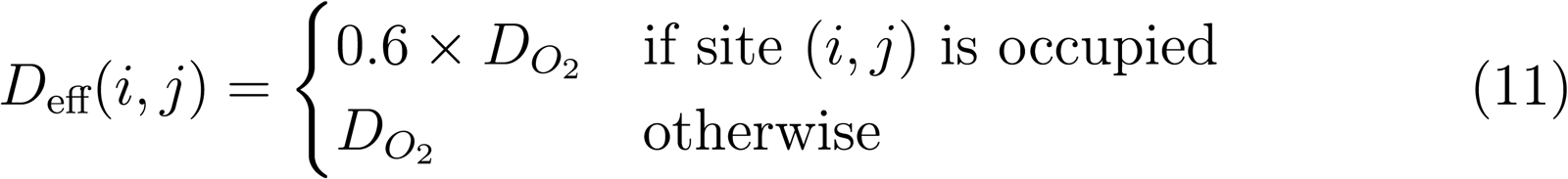

This modification reflects published estimates of mucin-mediated reduction in effective oxygen diffusivity in colorectal adenocarcinoma tissue of approximately 30–50% relative to normal tissue [33]. All other model parameters were held constant. A separate ensemble of 50 independent realisations was run under the 5*×*5 Gy SCRT protocol with this CRC-specific diffusion model to assess robustness of the primary findings.

## 3. Results

### 3.1. Spatial Oxygen Heterogeneity and the Hypoxic Niche

The initialised tumour lattice exhibited pronounced spatial oxygen heterogeneity consistent with diffusion-limited oxygen transport in avascular tumour tissue. At the pre-treatment timepoint, oxygen concentration ranged from normoxic values near vessel source sites to near-anoxic conditions in the tumour core, with a steep gradient across the peri-vascular to perinecrotic transition zone (Figure 1). ATP levels tracked the oxygen gradient through Michaelis–Menten kinetics, with cells in the hypoxic core operating exclusively on glycolytic metabolism. At the pre-treatment timepoint, a mean of 27 cells per realisation occupied the metabolic dormancy zone (6.4 *≤ ATP <* 16), representing the hypoxic niche population that is simultaneously protected from radiation by reduced OER and from immune killing by hypoxia-dependent CTL suppression. The local CTL kill probability map confirmed that immune killing capacity was near-zero in the hypoxic core and approached its maximum value exclusively in well-oxygenated perivascular regions, producing a spatial complementarity between the zones of radioresistance and immune evasion.

**Figure 1:**
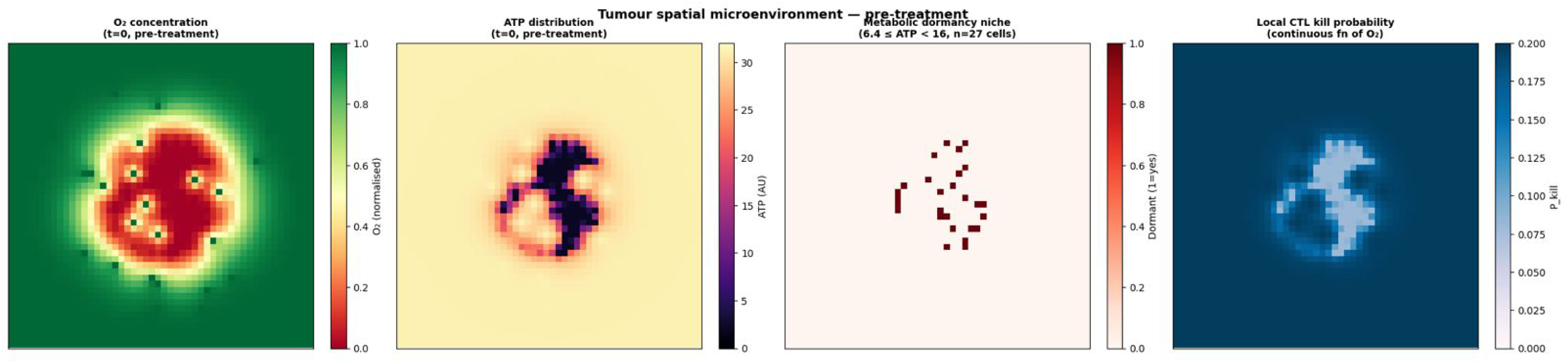
Tumour spatial microenvironment at the pre-treatment timepoint (*t* = 0). From left to right: oxygen concentration field (normalised, 0–1), with red indicating hypoxic regions and green vessel source sites; ATP distribution across tumour cells (AU), with dark purple indicating the metabolic dormancy zone (6.4 *≤ ATP <* 16); metabolic dormancy niche map showing the 27 cells occupying the dormancy zone at baseline; local CTL kill probability as a continuous function of *O*_2_, showing near-zero immune efficacy in the hypoxic core and maximum killing probability in peri-vascular regions. All panels from a single representative simulation run (RT+Immune, seed = 42).

To contextualise the modelled oxygen landscape against real patient biology, we quantified hypoxia burden across 284 Stage II/III CRC patients from TCGA-COAD and TCGA-READ using the Buffa 15-gene ssGSEA signature [32]. The resulting distribution was right-skewed (range 2.19–3.19, mean 2.63), with 57 patients (20.1%) classified as high-hypoxia above the 80th percentile threshold (Figure 2). This confirms that significant intratumoral hypoxia is a clinically prevalent feature of the target disease population, providing biological grounding for the oxygen heterogeneity incorporated in the model. We note that the Buffa score reflects bulk-tissue hypoxia averaged across all cell types, whereas the ABM quantifies cell-level oxygen at spatial resolution within the tumour boundary; direct numerical comparison is therefore not appropriate, but both measurements confirm that meaningful hypoxia is present in a substantial fraction of Stage II/III CRC.

**Figure 2:**
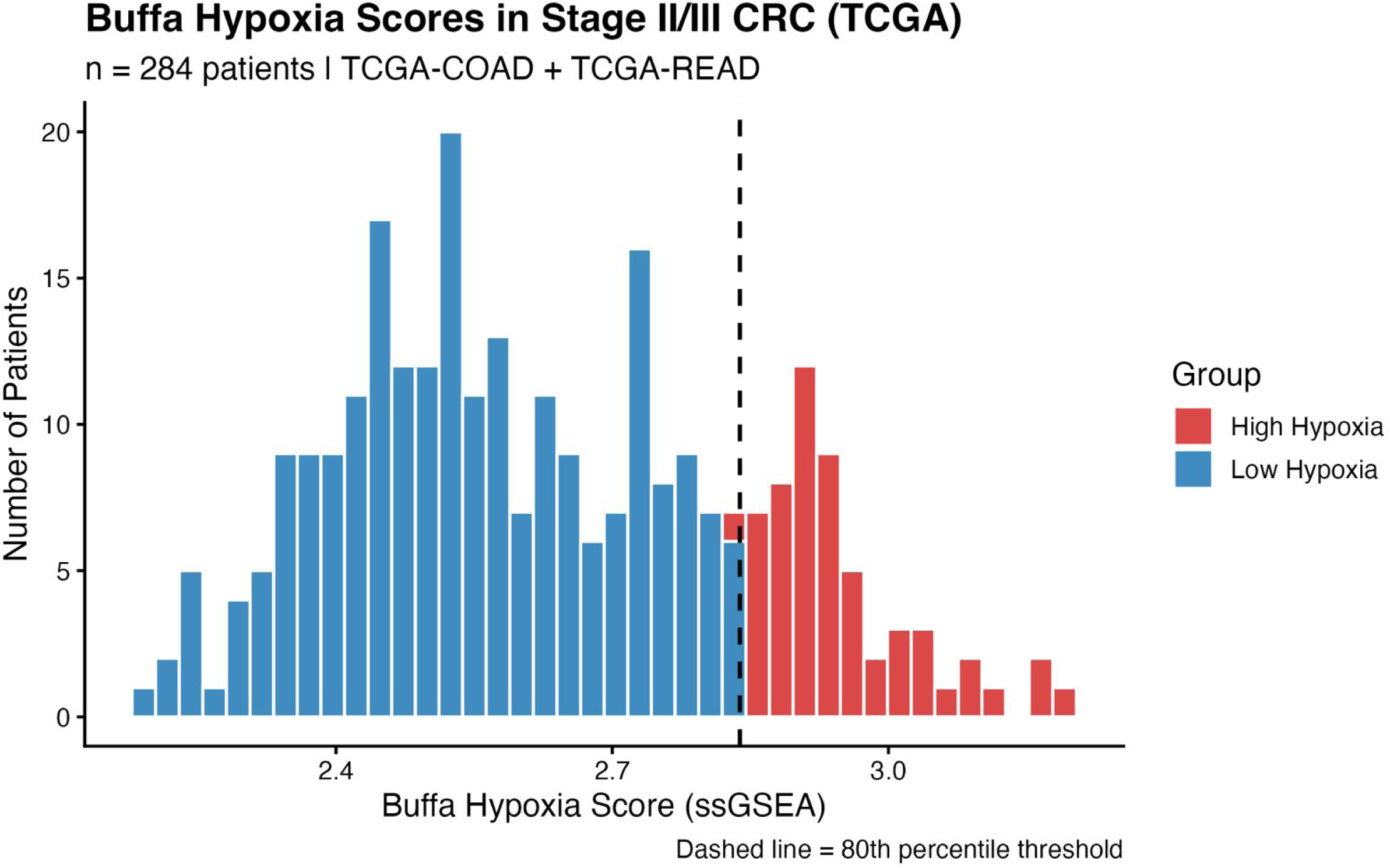
Tumour hypoxia burden in Stage II/III colorectal cancer from TCGA. Distribution of Buffa 15-gene ssGSEA hypoxia scores across 284 patients from TCGA-COAD and TCGA-READ with Stage II/III disease and matched RNA sequencing data. Blue bars: low-hypoxia patients (below 80th percentile). Red bars: high-hypoxia patients (above 80th percentile threshold, dashed vertical line), comprising 57 patients (20.1% of cohort). Score range: 2.19–3.19; mean: 2.63. The right-skewed distribution confirms that significant intratumoral hypoxia is a clinically prevalent but not universal feature of the target disease population, providing biological grounding for the oxygen heterogeneity modelled in this work.

### 3.2. Population Dynamics Under Fractionated Radiotherapy

Single-run trajectories revealed the characteristic sawtooth repopulation pattern expected from fractionated radiotherapy: each 5 Gy fraction produced a sharp reduction in live cell count, followed by partial recovery between fractions as surviving cells divided into vacated lattice space (Figure 3, Figure 4). This between-fraction repopulation, driven by resistant and hypoxia-protected cells expanding into space vacated by radiosensitive cells, is consistent with the accelerated repopulation phenomenon documented in clinical radiobiology [34]. The magnitude of repopulation between fractions was greater in the RT-only condition, where no immune clearance competed with cell division, than in the RT-plus-immune condition where CTL activity partially suppressed inter-fraction recovery.

**Figure 3:**
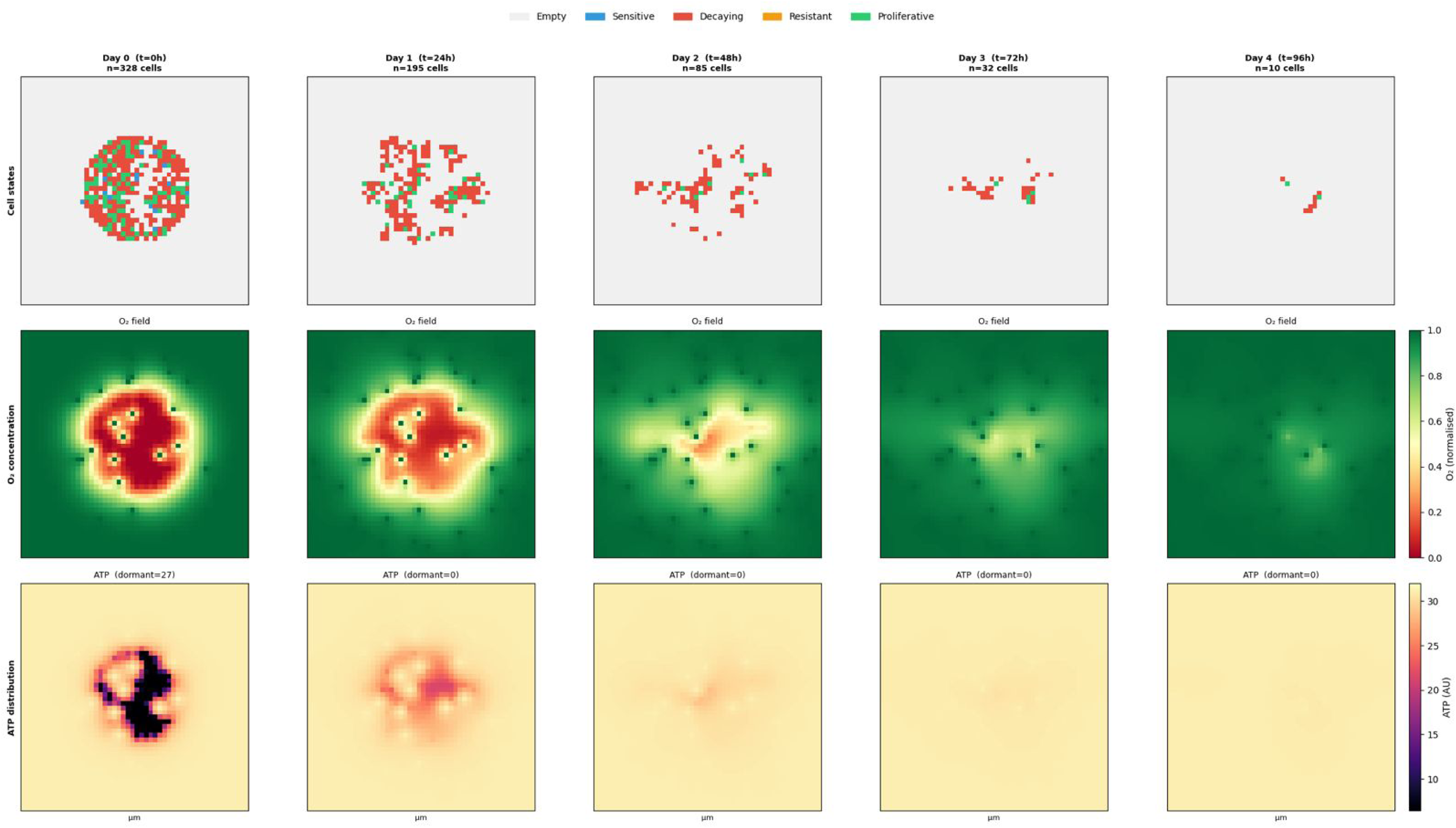
Spatial evolution of the tumour microenvironment across the radiotherapy course. Each column represents a timepoint corresponding to a radiotherapy fraction (Day 0 through Day 4, *t* = 0–96 h). Row 1: cell state maps showing the spatial distribution of sensitive (blue), proliferative (green), resistant (orange), and decaying (red) cells on the 50 *×* 50 lattice (1 mm2 tissue cross-section). Row 2: oxygen concentration field; vessel source sites appear as dark puncta against the normalised oxygen gradient. Row 3: ATP distribution across the lattice; subtitle indicates dormant cell count per timepoint (dormant = 27 at *t* = 0, falling to zero after fraction 1). All panels from a single representative simulation run (RT+Immune, seed = 42).

**Figure 4:**
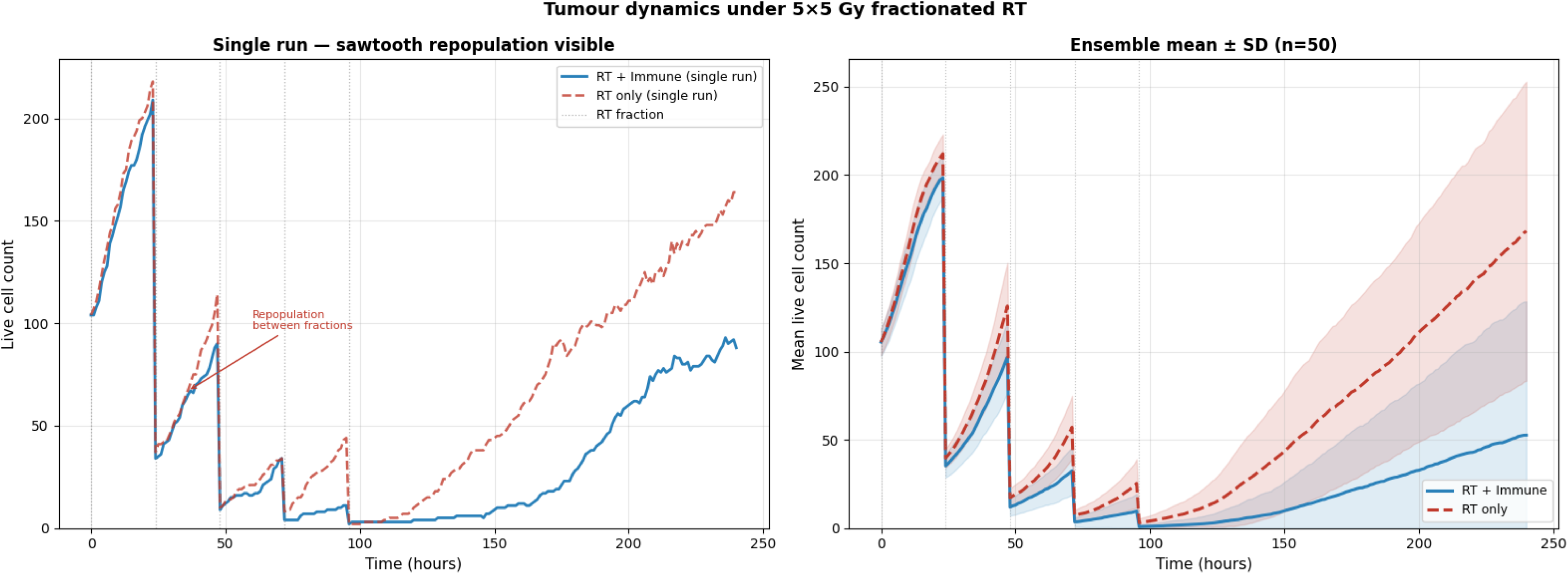
Tumour population dynamics under 5 *×* 5 Gy short-course radiotherapy. Left panel: single simulation run showing live cell count over time for RT+Immune (solid blue) and RT-only (dashed red) conditions. The sawtooth pattern reflects inter-fraction repopulation: sharp decline at each fraction followed by partial recovery as surviving cells divide into vacated lattice space. Annotated arrow indicates the repopulation phase. Right panel: ensemble mean *±* 1 standard deviation (shaded) across *n* = 50 independent realisations for each condition. Dotted vertical lines mark RT fraction timepoints.

Ensemble analysis (*n* = 50) confirmed that RT plus immune response produced significantly lower final tumour burden than RT alone (52.7 *±* 75.0 versus 168.2 *±* 83.8 cells at *t* = 240 h; Mann–Whitney *U*, *p* = 3.68 *×* 10*^−^*^9^) (Figure 5). The wide standard deviation in the RT-plus-immune condition reflects the bimodal outcome distribution: runs separated into a cleared group with near-zero final burden and an escaped group with substantial residual tumour, with minimal intermediate outcomes. Tumour control probability in the RT-plus-immune ensemble was 62.0% (95% Wilson CI: 0.482–0.741), compared to 0% in the RT-only ensemble by the same criterion (final burden*<* 5 cells).

**Figure 5:**
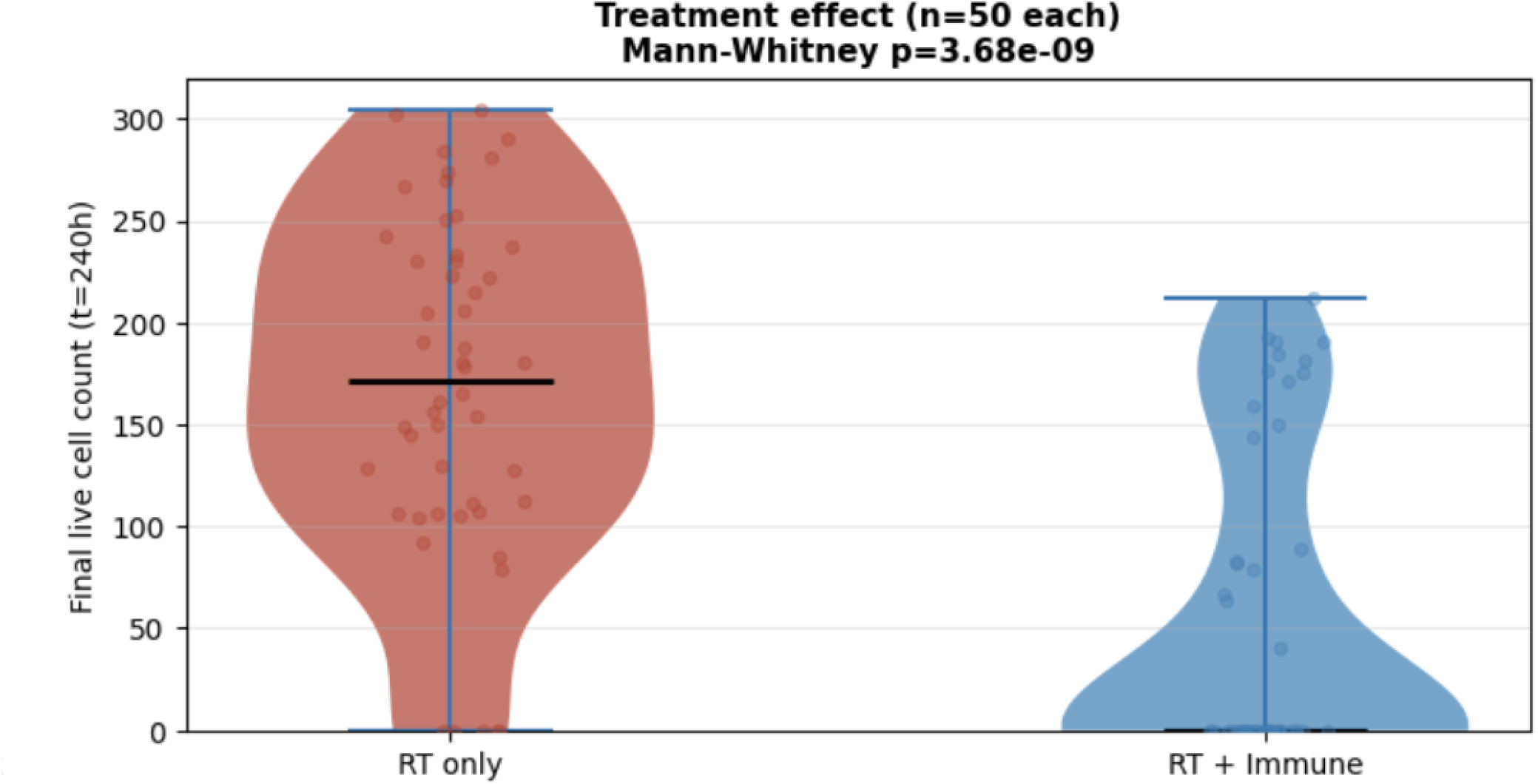
Final tumour burden at *t* = 240 h for RT-only and RT+Immune ensembles (*n* = 50 each). Violin plots show the full distribution of final live cell counts; individual run values are overlaid as scatter points; horizontal bars indicate the median. The RT+Immune condition produces a bimodal distribution reflecting cure-or-escape outcomes, in contrast to the unimodal high-burden distribution under RT alone. Mann–Whitney *U* test: *p* = 3.68 *×* 10*^−^*^9^.

### 3.3. Clonal Evolution and Subpopulation Dynamics

Subpopulation analysis revealed progressive enrichment of the resistant clone over the treatment course, consistent with Darwinian selection under cytotoxic pressure (Figure 6). Each radiation fraction preferentially eliminated sensitive and proliferative cells, which comprised the majority of the tumour mass and carried the highest radiosensitivity parameters. The declining kill per fraction—from a mean of 270 cells eliminated in fraction 1 to 42 cells in fraction 5—quantified this progressive resistant enrichment directly (Figure 7). By fraction 5, the surviving population was substantially depleted of radiosensitive cells, explaining the marked reduction in per-fraction cytotoxicity over the treatment course.

**Figure 6:**
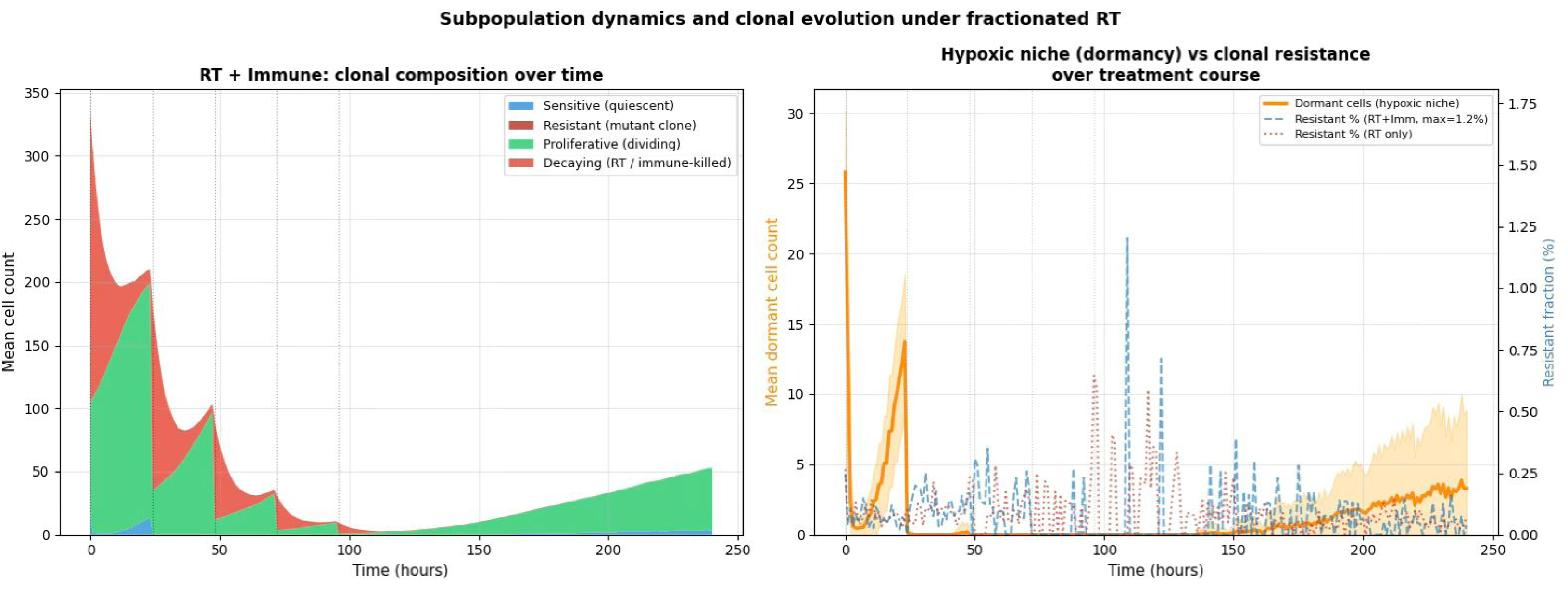
Subpopulation composition and hypoxic niche dynamics over the treatment course. Left panel: stackplot showing mean cell count per subpopulation (sensitive, resistant, proliferative, decaying) across time in the RT+Immune ensemble (*n* = 50). Dotted vertical lines mark RT fraction timepoints. Right panel: mean dormant cell count (orange, left axis, *±*1 SD shaded) and resistant clone fraction as percentage of total live cells (dashed lines, right axis) over the treatment course, for RT+Immune (blue dashed) and RT-only (red dotted) ensembles. The resistant fraction maximum (1.2%) is indicated in the legend.

**Figure 7:**
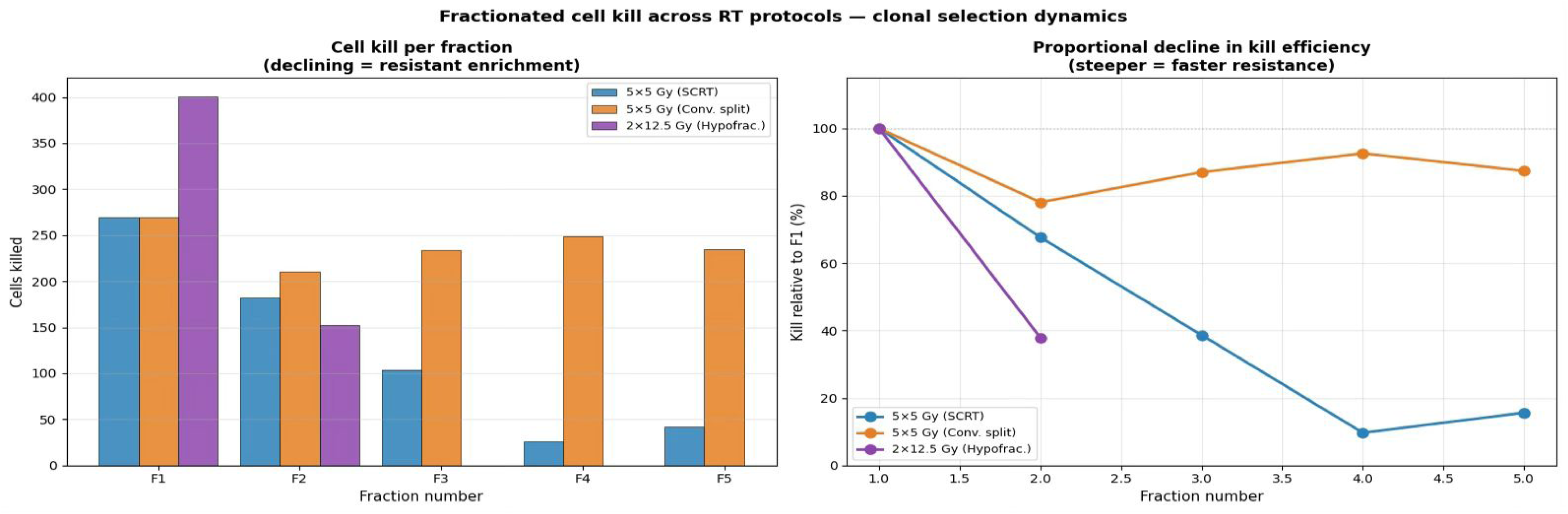
Fractionated cell kill across radiotherapy protocols. Left panel: absolute number of cells killed per fraction for each protocol in a single representative RT-only run (immune response disabled to isolate radiation-specific kill). Right panel: cell kill per fraction expressed as a percentage of fraction-1 kill, showing the proportional decline in kill efficiency across fractions. A steeper decline indicates faster enrichment of the radioresistant subpopulation. 5 *×* 5 Gy SCRT (blue), 5 *×* 5 Gy conventional split (orange), 2 *×* 12.5 Gy hypofractionation (purple).

The resistant clone, however, never exceeded 1.2% of the total tumour population at any timepoint in either ensemble. This finding indicates that the dominant escape mechanism in the model is not mutational resistance but rather spatial protection conferred by the hypoxic microenvironment—cells surviving in the hypoxic niche retain sensitive phenotypes but are shielded from both radiation and immune killing by their local oxygen environment. The dormancy reservoir, comprising cells in the metabolic dormancy zone, peaked transiently at the pre-treatment timepoint (mean 27 cells) and appeared briefly during inter-fraction intervals before being progressively eliminated as the tumour regressed and oxygen redistributed into the formerly hypoxic core.

### 3.4. Single-Cell Mechanistic Trajectories

Agent-level tracking revealed five qualitatively distinct cell fate categories that together illustrate the mechanistic basis of both treatment success and failure (Figure 8). Cells killed in the first fraction occupied intermediate-oxygenation positions and carried sensitive-phenotype radiosensitivity parameters, dying within 25 hours of the initial dose. Persistently dormant cells maintained oxygen levels near zero throughout the simulation, with ATP fluctuating in the dormant zone (6.4–16 AU) on pure glycolytic support—alive but energetically unable to divide, and invisible to immune killing given nearzero local CTL kill probability. The dormant-to-escape trajectory demonstrated the wake-up mechanism central to the paper’s biological argument: a cell beginning in the hypoxic core experienced progressive oxygen improvement as surrounding cells died and oxygen redistributed, crossing the ATP proliferation threshold and resuming division in the post-treatment window. Resistant clone trajectories showed sustained survival through all five fractions at intermediate oxygen levels, with radiosensitivity too low for the linear-quadratic survival probability to produce death even under normoxic conditions. Finally, immune-cleared survivors demonstrated the spatial vulnerability of normoxic survivors: cells that escaped radiation by chance found themselves in well-oxygenated regions where CTL density accumulated and killing probability was maximal, resulting in post-treatment elimination despite having survived all five fractions.

**Figure 8:**
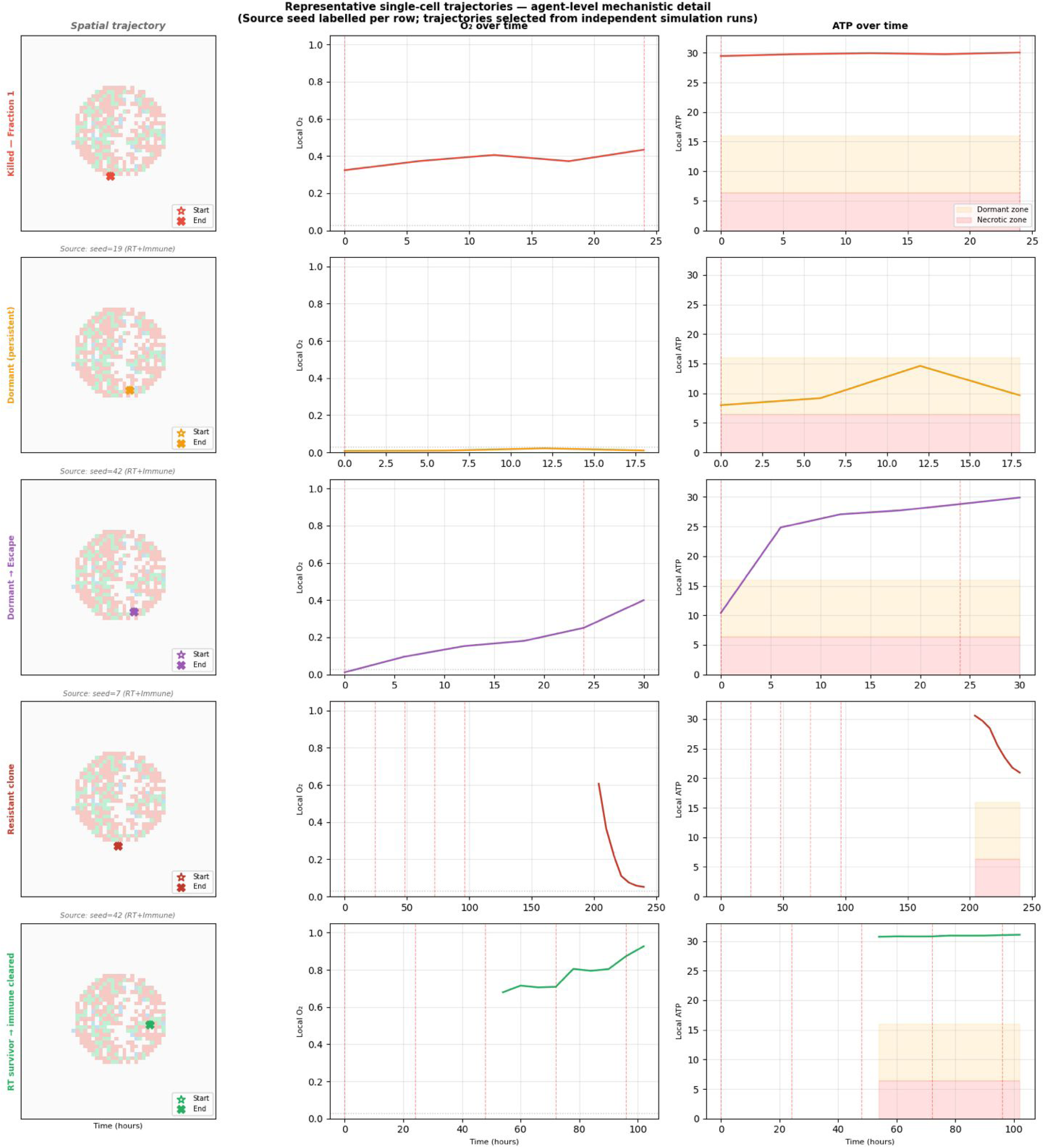
Representative single-cell trajectories illustrating agent-level mechanistic detail. Five fate categories are shown, one per row: cells killed in the first RT fraction (red), persistently dormant cells (orange), cells transitioning from dormancy to post-treatment escape (purple), resistant clones surviving all fractions (dark red), and RT survivors subsequently cleared by immune killing (green). For each cell, three panels are shown: spatial trajectory overlaid on the pre-treatment tumour grid (left column), local *O*_2_ over time (centre column), and local ATP over time with dormancy zone (orange band, 6.4–16 AU) and necrotic zone (red band, *<* 6.4 AU) indicated (right column). Red dashed vertical lines mark RT fraction timepoints. Source simulation seed is labelled above each spatial trajectory panel; representative cells were identified by searching across independent simulation runs.

**Figure 9:**
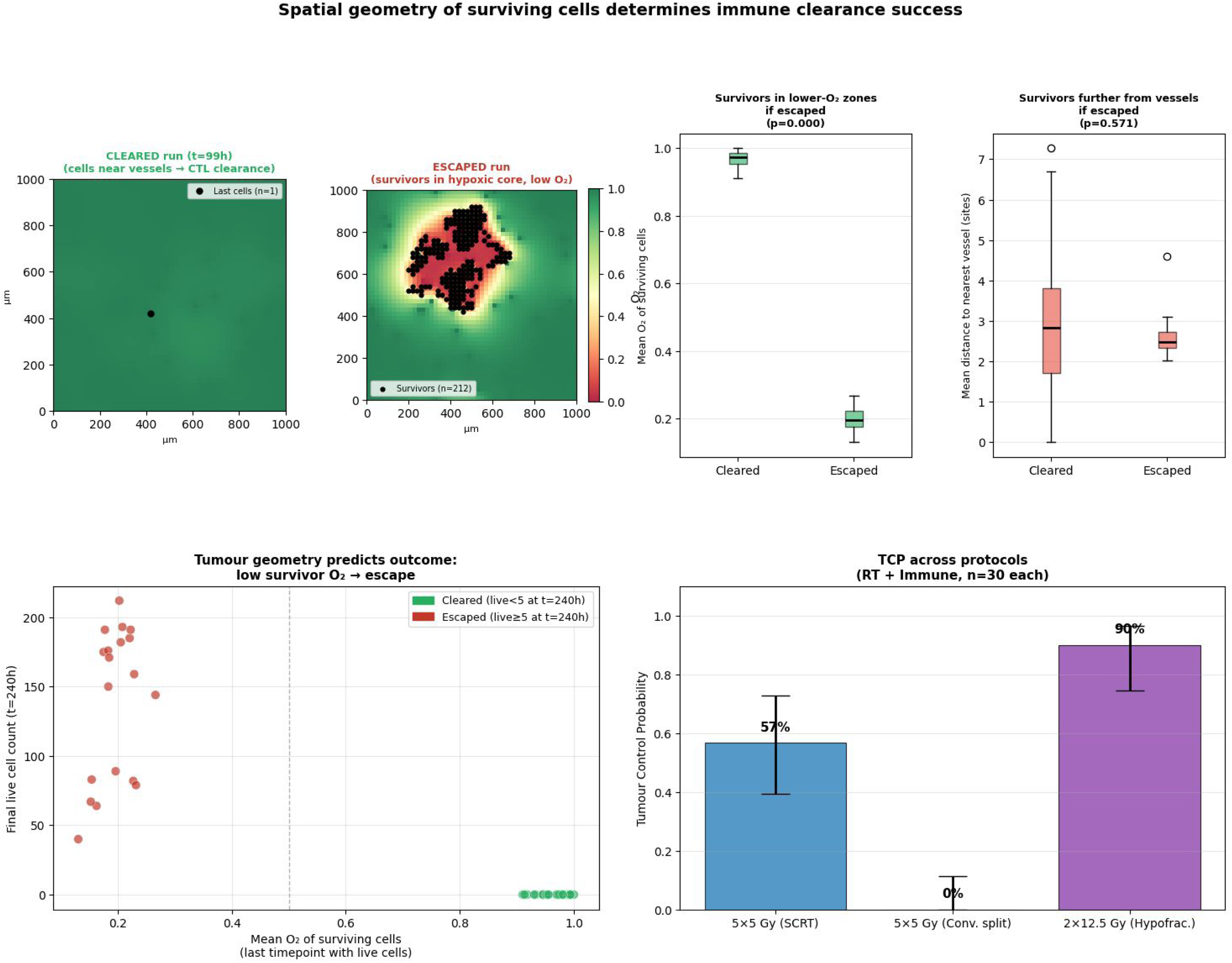
Spatial geometry of surviving cells determines immune clearance success. Top row, left: spatial map of a cleared run at the last timepoint with live cells present, showing the *O*_2_ field with surviving cell positions (black dots); cells are located in well-oxygenated regions adjacent to vessel sites. Top row, centre: spatial map of an escaped run (*t* = 240 h) showing surviving cells concentrated in the hypoxic core. Top row, right two panels: boxplots comparing mean *O*_2_ of surviving cells (left) and mean distance to nearest vessel (right) between cleared (*n* = 31) and escaped (*n* = 19) runs; *p*-values from Mann–Whitney *U* test indicated. Bottom row, left: scatter plot of final tumour burden versus mean *O*_2_ of surviving cells at the last timepoint with live cells, for all 50 realisations; cleared runs (green) and escaped runs (red) are completely separated along the *O*_2_ axis with no overlap. Bottom row, right: tumour control probability across three equal total-dose protocols (*n* = 30 per arm), with Wilson 95% confidence intervals.

### 3.5. Spatial Geometry of the Hypoxic Niche Predicts Immune Clearance

Of 50 ensemble realisations, 31 achieved tumour control (final burden *<* 5 cells) and 19 experienced treatment escape with sustained tumour regrowth. Spatial analysis of surviving cell positions at the last timepoint with live cells present revealed a striking difference in microenvironmental context between the two outcome groups (Figure 10).

**Figure 10:**
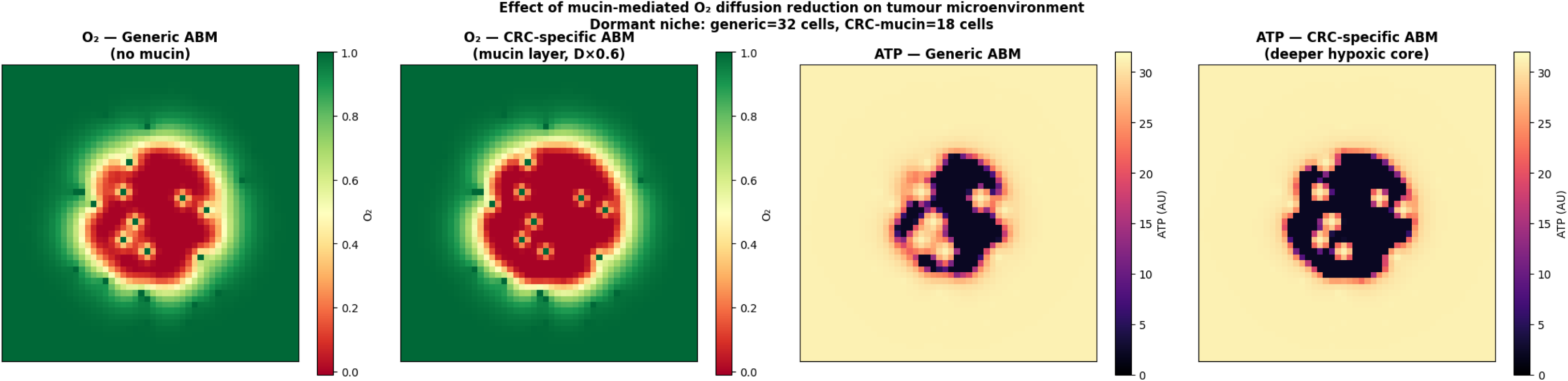
Effect of mucin-mediated oxygen diffusion reduction on the CRC tumour microenvironment. Four spatial maps comparing the generic ABM (left two panels) and the CRC-specific mucin model (right two panels) at the pre-treatment timepoint: O concentration and ATP distribution. The CRC-mucin model produces a deeper hypoxic core and larger necrotic zone consistent with mucin-secreting colorectal adenocarcinoma tissue physiology. Dormant niche cell count: generic = 32 cells, CRC-mucin = 18 cells.

In cleared runs, the final surviving cells occupied well-oxygenated regions adjacent to vessel source sites, with mean local *O*_2_ = 0.967 across the cleared group. In escaped runs, surviving cells were concentrated in the hypoxic core, with mean local *O*_2_ = 0.196. This difference was highly significant (Mann–Whitney *U*, *p* = 2.1 *×* 10*^−^*^9^) and was consistent across all 50 realisations without exception: no escaped run had mean survivor *O*_2_ above 0.27, and no cleared run had mean survivor *O*_2_ below 0.91, producing complete separation between outcome groups with no overlap across all 50 realisations. The scatter of final tumour burden against mean survivor *O*_2_ demonstrated complete separation between outcome groups along the oxygen axis, with escaped tumours clustering between *O*_2_ = 0.10–0.35 and cleared tumours clustering between *O*_2_ = 0.85–1.00.

Mean distance from surviving cells to the nearest vessel source site did not differ significantly between cleared and escaped runs (2.91 versus 2.62 lattice sites; Mann–Whitney *U*, *p* = 0.571). This null result is mechanistically informative: vessel proximity is an imperfect proxy for local oxygenation because cellular oxygen consumption creates depletion gradients that can render vessel-adjacent sites hypoxic under high tumour cell density. Direct measurement of local *O*_2_ is therefore a more precise predictor of immune clearance outcome than anatomical distance to vasculature, supporting the use of the continuous oxygen field as the primary microenvironmental readout in this model.

These findings support the hypothesis that spatial position within the oxygen landscape—rather than aggregate surviving cell count—is a primary determinant of whether CTL-mediated clearance succeeds following fractionated radiotherapy. The mechanism is the spatial coincidence of two independent protection gradients: radiation efficacy decreases and immune killing probability decrease by the same continuous function of local *O*_2_, creating a microenvironmental refuge in which cells surviving radiation are simultaneously the least accessible to immune clearance.

### 3.6. Robustness of spatial findings under CRC-specific microenvironmental conditions

To verify that the central finding of this study is not an artefact of generic tumour model assumptions, we assessed the effect of mucin-mediated oxygen diffusion reduction on treatment outcome and spatial oxygen geometry. The CRC-specific model produced a visibly deeper hypoxic core relative to the generic model, with a larger necrotic zone and altered dormancy niche geometry at the pre-treatment timepoint (generic: 32 dormant cells; CRC-mucin: 18 dormant cells), consistent with the more severe intratumoral hypoxia expected in mucin-secreting colorectal adenocarcinoma tissue.

Despite this CRC-specific modification to the oxygen microenvironment, tumour control probability under the 5*×*5 Gy SCRT protocol with concurrent immune response was unchanged at 62.0% (95% CI: [0.482, 0.741]; *n* = 50), identical to the primary ensemble. The difference in final tumour burden between the CRC-specific and generic models was not statistically significant (Mann–Whitney *U*, *p* = 0.293). The spatial separation in survivor oxygen concentration between cleared and escaped runs was preserved under the mucin model, confirming that the relationship between oxygen microenvironment geometry and immune clearance outcome is robust to CRC-specific tissue diffusion properties. These results indicate that the primary finding reflects a general property of spatially heterogeneous oxygen landscapes under combined RT-immune treatment, grounded in CRC-relevant microenvironmental parameters.

### 3.7. Global Sensitivity Analysis

Sensitivity analysis across four parameters confirmed that radiosensitivity (*α*) was the dominant determinant of final tumour burden (Pearson *r* = *−*0.629), followed by base proliferation rate (*r* = +0.402), maximum CTL kill probability (*r* = *−*0.109), and oxygen half-saturation constant *K_m_*(*r* = +0.085) (Figure 11). The strong negative correlation with *α* confirms that the model’s primary cytotoxic mechanism—radiation-induced cell death—behaves as expected: higher radiosensitivity produces lower residual burden. The positive correlation with proliferation rate reflects the competition between treatment-induced killing and between-fraction repopulation. The weak correlations with immune killing strength and metabolic parameters indicate that model outcomes are not dominated by immune parameter choices, supporting the robustness of the spatial clustering finding to uncertainty in CTL efficacy. These rankings are consistent with established clinical understanding of CRC radiotherapy, in which tumour radiosensitivity is a dominant determinant of pathological complete response [35, 13].

**Figure 11:**
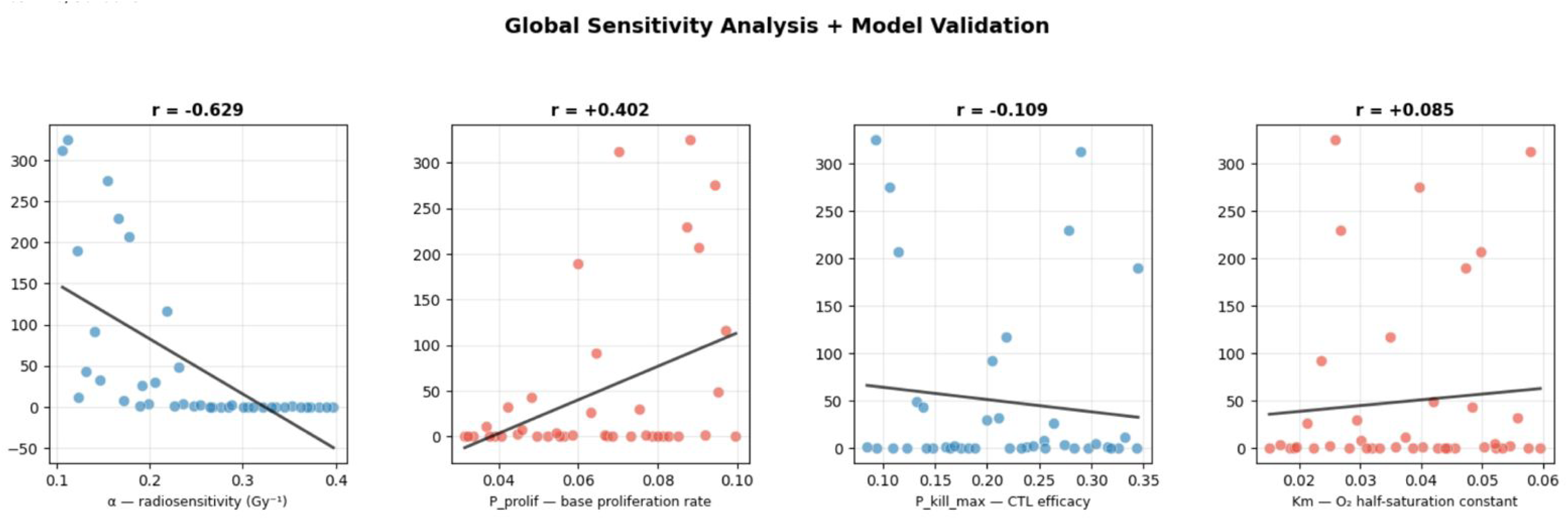
Global sensitivity analysis. Scatter plots of final tumour burden against each of four parameters varied by Latin Hypercube Sampling (*n* = 40 samples, 3 replicates each): radiosensitivity *α* (*r* = *−*0.629), base proliferation rate (*r* = +0.402), maximum CTL kill probability (*r* = *−*0.109), and *O*_2_ half-saturation constant *K_m_* (*r* = +0.085). Blue points: cleared runs (final burden *<* 5 cells). Red points: escaped runs. Trend lines from linear regression shown in black.

**Figure 12:**
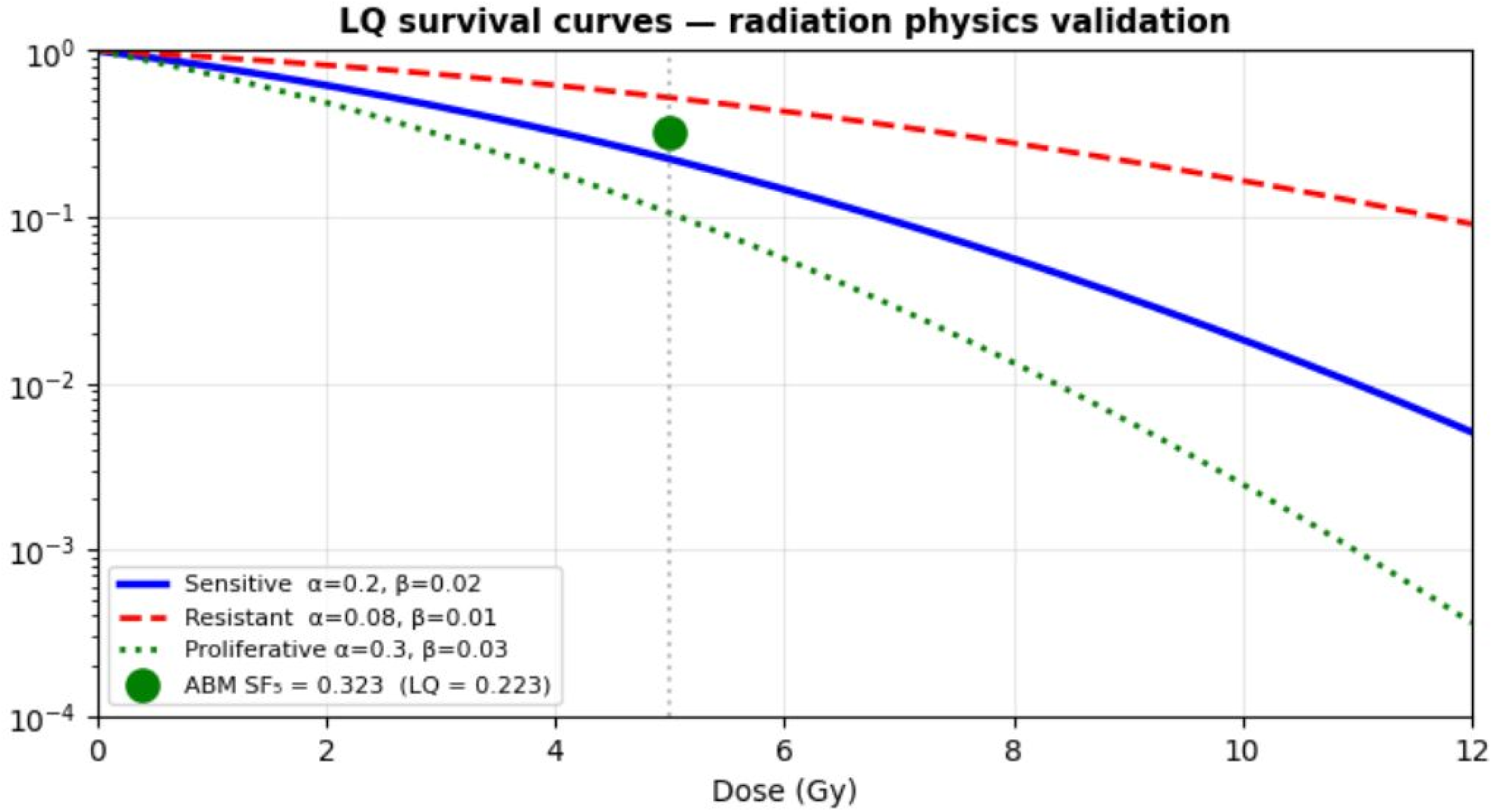
Linear-quadratic survival curves for the three cell phenotypes implemented in the model: sensitive cells (*α* = 0.20 Gy*−*1, *β* = 0.02 Gy*−*2, solid blue), resistant cells (*α* = 0.08 Gy*−*1, *β* = 0.01 Gy*−*2, dashed red), and proliferative cells (*α* = 0.30 Gy*−*1, *β* = 0.03 Gy*−*2, dotted green). The ABM-observed population-averaged survival fraction following a single 5 Gy fraction (*SF*_5_ = 0.323, green circle) lies above the normoxic sensitive-cell LQ prediction (*SF*_5_ = 0.223, dotted vertical line); the offset is attributable to the radioresistant hypoxic subpopulation included in the spatial average, confirming internal consistency of the radiation physics implementation.

### 3.8. Fractionation Schedule and Tumour Control Probability

Protocol comparison across three equal total-dose schedules (25 Gy each) demonstrated that inter-fraction interval is a critical determinant of tumour control probability, independent of cumulative dose (Figure 13, Figure 7).

**Figure 13:**
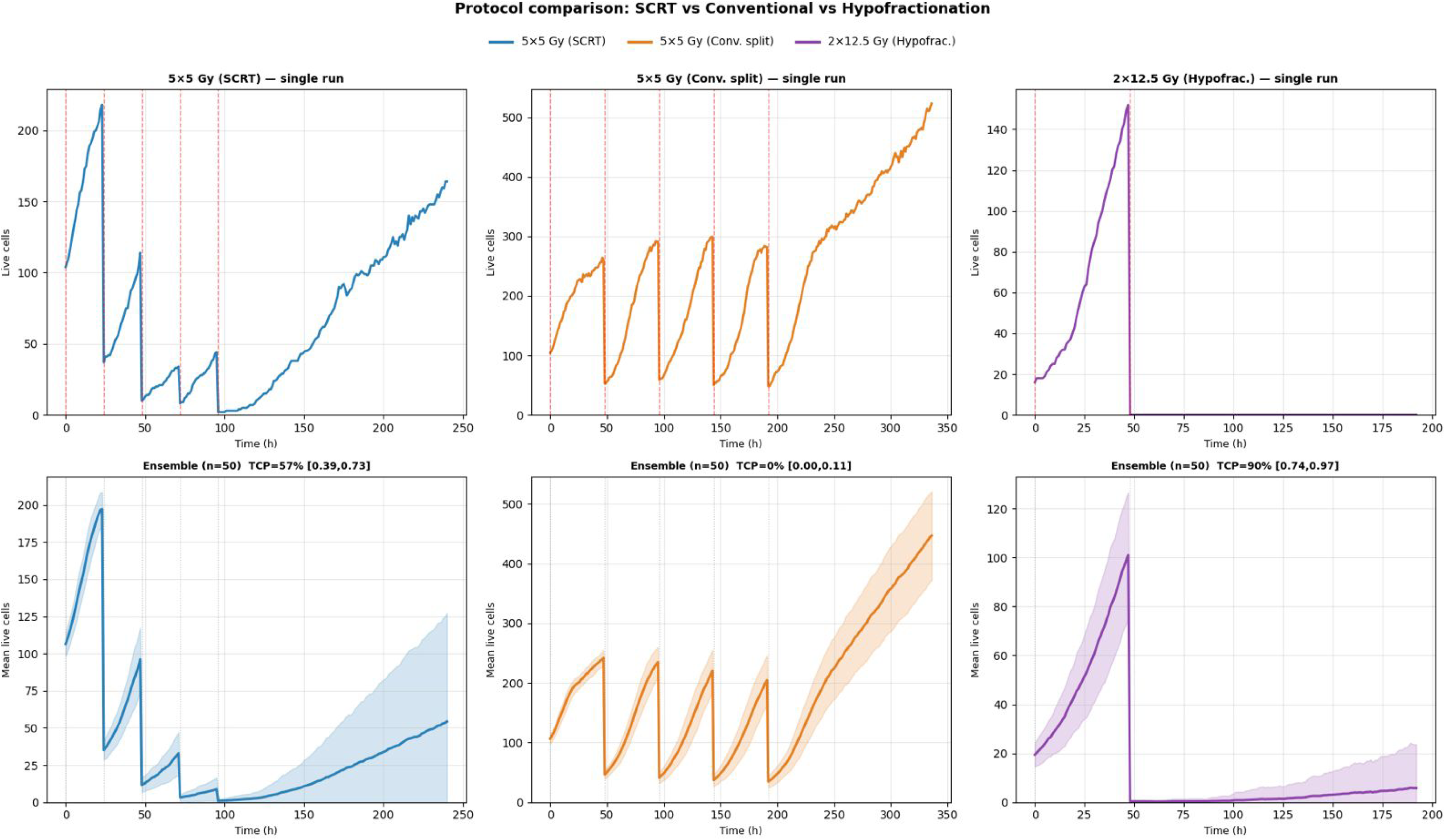
Protocol comparison: tumour dynamics across three equal total-dose schedules. Three radiotherapy protocols each delivering 25 Gy total are compared: 5 *×* 5 Gy SCRT (blue, daily fractions), 5 *×* 5 Gy wide spacing (orange, 48-hour inter-fraction intervals), and 2 *×* 12.5 Gy hypofractionation (purple, single 48-hour interval). Top row: single representative simulation run per protocol, showing live cell count over time. Bottom row: ensemble mean *±* 1 standard deviation (*n* = 30 per arm), with tumour control probability and Wilson 95% confidence interval indicated in panel title. Red dashed vertical lines mark RT fraction timepoints. All protocols include concurrent immune response.

The standard SCRT protocol (5 *×* 5 Gy, daily fractions) achieved TCP = 57% (95% CI: 0.39–0.73) with mean final burden of 54 *±* 72 cells. The widely-spaced fractionation schedule (5 *×* 5 Gy, 48-hour inter-fraction intervals) achieved TCP = 0% (95% CI: 0.00–0.11) with mean final burden of 446 *±* 73 cells, as extended inter-fraction intervals permitted substantial tumour repopulation that overwhelmed per-fraction cytotoxicity. Hypofractionation (2 *×* 12.5 Gy, single 48-hour interval) achieved the highest TCP of 90% (95% CI: 0.74–0.97) with mean final burden of 6 *±* 18 cells, consistent with the minimal repopulation window afforded by two large fractions.

Cell kill efficiency declined across fractions in all protocols, but the rate of decline differed markedly by schedule (Figure 7). The SCRT protocol showed a progressive decline from fraction 1 to fraction 5, reflecting resistant clone enrichment over daily fractionation. The widely-spaced protocol maintained relatively stable per-fraction kill across fractions because extended intervals allowed sensitive cell repopulation, presenting each fraction with a partially refreshed sensitive population—but this repopulation simultaneously drove tumour growth between fractions, producing net treatment failure. These findings suggest that inter-fraction interval exerts opposing effects on per-fraction kill efficiency and cumulative tumour control, with compact scheduling favouring overall control despite accelerated resistant enrichment within the treatment course. We note that immune parameters were held constant across protocols as a deliberate methodological choice to isolate fractionation schedule effects; the differential immunogenic cell death kinetics that may accompany different fractionation schedules in vivo represent an important direction for future investigation.

**Table 2:** Tumour control probability and mean final tumour burden across three equal total-dose radiotherapy protocols (25 Gy each), all with concurrent immune response (n = 30 realisations per arm). TCP defined as fraction of realisations with final live cell count below 5. Confidence intervals computed using the Wilson score method. Mean burden reported as mean ± standard deviation at the final simulation timepoint for each protocol.

| **Protocol** | **Total dose** | **Fractions** | **TCP** | **95% CI** | **Mean final burden (cells)** |
|---|---|---|---|---|---|
| 5 × 5 Gy (SCRT) | 25 Gy | 5 × daily | 57% | [0.39, 0.73] | 54 ± 72 |
| 5 × 5 Gy (wide spacing) | 25 Gy | 5 × 48 h | 0% | [0.00, 0.11] | 446 ± 73 |
| 2 × 12.5 Gy (hypofrac.) | 25 Gy | 2 × 48 h | 90% | [0.74, 0.97] | 6 ± 18 |

## 4. Discussion

### 4.1. Spatial Oxygen Geometry as a Determinant of Treatment Outcome

The central finding of this study is that the spatial position of surviving cells within the tumour oxygen landscape at the end of radiotherapy—not the number of surviving cells alone—predicts whether cytotoxic T lymphocytemediated clearance succeeds or fails. Across 50 independent realisations of the same treatment protocol, outcome was determined by microenvironmental context with complete separation: cleared tumours retained their final surviving cells in well-oxygenated regions (mean *O*_2_ = 0.967, minimum 0.910), while escaped tumours had surviving cells concentrated in the hypoxic core (mean *O*_2_ = 0.196, maximum 0.266), with no overlap between groups across any realisation. The mechanistic basis of this separation is the spatial coincidence of two independent protection gradients that share the same functional dependence on local oxygen: radiation efficacy, governed by the oxygen enhancement ratio, and immune killing probability, governed by CTL effector function under hypoxia. Where one fails, the other fails by the same physics. The hypoxic niche is not simply radioresistant—it is simultaneously immune-evasive, and this double protection is what the model identifies as the functional determinant of treatment failure.

This finding is consistent with a growing body of experimental and clinical evidence linking intratumoral hypoxia to both radiotherapy resistance and immune exclusion in colorectal cancer [13, 15, 16, 36, 11, 12]. What the present model contributes is a spatially explicit mechanistic account of how these two phenomena interact at the level of individual cells and their microenvironmental context, producing an emergent population-level outcome—the bimodal cure-or-escape distribution—that neither a purely radiobiological nor a purely immunological model could generate in isolation. The TCGA transcriptomic analysis confirmed that significant hypoxia is a clinically prevalent feature of Stage II/III CRC, with 20.1% of 284 patients classified as high-hypoxia by the Buffa ssGSEA signature [32], providing biological grounding for the oxygen heterogeneity incorporated in the model.

### 4.2. The Hypoxic Niche as Microenvironmental Refuge

The metabolic dormancy zone identified in this model—cells sustaining viability through anaerobic glycolysis at ATP levels below the proliferation threshold but above the necrotic threshold—represents a transient but mechanistically significant microenvironmental state. Dormant cells peaked at a mean of 27 per realisation at the pre-treatment timepoint and appeared briefly during inter-fraction intervals, before being progressively eliminated as tumour regression allowed oxygen redistribution into formerly hypoxic regions. This behaviour is consistent with the peri-necrotic hypoxic gradient described in solid tumour physiology, in which cells at intermediate distances from functional vasculature occupy a metabolic niche characterised by chronic oxygen deficiency insufficient to drive immediate necrosis but sufficient to impair proliferative capacity [10, 37].

We deliberately avoid framing dormancy as the primary determinant of radiotherapy failure in this model. The data show that the dominant escape mechanism is spatial *O*_2_-dependent protection of cells that retain their sensitive phenotype—not metabolic dormancy per se, and not mutational resistance, which never exceeded 1.2% of the tumour population. The hypoxic niche is better characterised as a microenvironmental refuge that functions through the coincident suppression of two cytotoxic mechanisms, rather than as a stable dormant reservoir driving long-term recurrence. This distinction matters for clinical translation: it suggests that strategies targeting the oxygen microenvironment—reoxygenation, anti-angiogenic normalisation, or hypoxia-activated prodrugs—may be more effective than strategies targeting dormancy pathways alone, because the protection is conferred by the environment rather than by an intrinsic cellular state [38].

### 4.3. Fractionation Schedule and Immune–RT Timing

The protocol comparison demonstrated that inter-fraction interval is a critical modulator of tumour control probability independent of cumulative dose. The widely-spaced fractionation schedule—identical total dose to SCRT but delivered over twice the calendar time—achieved zero tumour control across 30 realisations, as extended inter-fraction intervals permitted tumour repopulation that overwhelmed cumulative cytotoxicity. This result is consistent with the overall treatment time effect documented in clinical radiobiology, in which prolongation of the treatment course beyond the accelerated repopulation threshold substantially reduces locoregional control [39, 34].

The hypofractionation protocol (2*×*12.5 Gy) achieved the highest TCP of 90%, suggesting that compressing equivalent dose into fewer, larger fractions minimises the repopulation window and maximises cumulative cytotoxicity. This finding aligns with the growing clinical evidence base for hypofractionation in rectal cancer, in which short-course protocols have demonstrated non-inferior or superior locoregional control compared to prolonged chemoradiotherapy schedules in several randomised trials [3, 4]. Mechanistically, the model predicts that hypofractionation’s advantage arises not from higher percell kill alone—the linear-quadratic model accounts for this through the *β* term—but from the reduced opportunity for between-fraction repopulation by hypoxia-protected and resistant subpopulations.

We note an important caveat. Immune parameters were held constant across all three protocols as a deliberate methodological choice to isolate fractionation schedule effects from immune response differences. In biological reality, different fractionation schedules may elicit distinct immunogenic cell death kinetics, with evidence suggesting that intermediate doses per fraction (around 8–12 Gy) may be optimal for stimulating abscopal immune responses [6, 40]. Exploring immune parameter sensitivity across fractionation schedules represents an important direction for future computational and experimental investigation. The present results should therefore be interpreted as isolating the repopulation-driven component of fractionation schedule effects, with immune–RT interaction dynamics providing an additional layer of complexity beyond the current model scope.

### 4.4. Model Validation and Parameter Sensitivity

The radiation physics implementation was validated by comparing the population-averaged survival fraction following a single 5 Gy fraction (*SF*_5_ = 0.323) against the normoxic linear-quadratic prediction for sensitive cells (*SF*_5_ = 0.223). The 44.6% difference reflects the spatially averaged response across the full oxygen distribution, in which the hypoxic subpopulation contributes disproportionately to overall survival by virtue of reduced OER. This is an expected and mechanistically interpretable discrepancy rather than a calibration error: a spatially homogeneous model with uniform normoxic parameters would reproduce the linear-quadratic prediction exactly, but would eliminate the hypoxic heterogeneity that is the primary biological question of this work. The *SF*_5_ gap is therefore evidence that the spatial oxygen model is functioning as intended, not evidence of miscalibration.

Global sensitivity analysis confirmed that model outcomes are dominated by biologically appropriate parameters. Radiosensitivity (*α*, *r* = *−*0.629) was the strongest determinant of final tumour burden, consistent with its established role as a primary predictor of pathological complete response in clinical CRC radiotherapy [35, 13]. Proliferation rate (*r* = +0.402) was the second strongest, reflecting the competition between cytotoxicity and between-fraction repopulation. CTL killing strength (*r* = *−*0.109) and the oxygen half-saturation constant *K_m_* (*r* = +0.085) showed weak correlations, indicating that the spatial clustering finding is not an artefact of specific immune or metabolic parameter choices but emerges robustly across the parameter space explored. Ensemble convergence was confirmed by bootstrap resampling, with TCP and burden estimates stabilising at *n ≥* 25 and 90% confidence interval width falling below 10% at *n* = 45.

The distance-to-vessel metric did not significantly differ between cleared and escaped runs (*p* = 0.571), despite the highly significant difference in local *O*_2_ (*p* = 2.1 *×* 10*^−^*^9^). This dissociation reflects a known feature of oxygen transport in tumour tissue: cellular oxygen consumption creates depletion gradients that render vessel-adjacent sites hypoxic under conditions of high tumour cell density, decoupling anatomical vessel proximity from functional oxygenation [12]. This finding reinforces the importance of measuring the oxygen field directly rather than using vascular geometry as a proxy, and supports the mechanistic design choice of computing *O*_2_ at each lattice site through explicit diffusion rather than estimating it from vessel distance alone.

### 4.5. Limitations and Future Directions

Several limitations of the present model should be acknowledged. The lattice geometry is two-dimensional, representing a tissue cross-section rather than a volumetric tumour mass. While two-dimensional ABMs capture the essential physics of oxygen diffusion and spatial cellular interaction relevant to the biological questions addressed here, and have been extensively used in computational oncology for this purpose [41, 22, 42, 29], extension to three dimensions would more accurately represent the full spatial complexity of tumour oxygenation and immune infiltration. Vascular geometry was represented by fixed point sources at 2% lattice density, without dynamic remodelling in response to treatment; incorporating vascular normalisation or RT-induced vascular damage would add biological realism to the oxygen dynamics. The model does not incorporate explicit genomic instability beyond the stochastic single-locus resistance mutation, and does not represent stromal or immune cell populations beyond the CTL field; integrating cancerassociated fibroblast interactions and regulatory T cell dynamics would extend the immune module toward greater biological completeness.

The *SF*_5_ discrepancy between the ABM (0.323) and the normoxic linear-quadratic prediction (0.223) warrants further investigation. Future work should compare modelled *pO*_2_ distributions against direct needle electrode measurements in CRC tissue to constrain the oxygen parameter space, following the approach of Vaupel and colleagues [10, 37]. Additionally, the empirical *O*_2_ threshold identified in this model—below which immune clearance never succeeded (*O*_2_ *<* 0.27) and above which it always succeeded (*O*_2_ *>* 0.91)—represents a testable prediction that could motivate prospective stratification of CRC patients by intratumoral hypoxia burden prior to SCRT. Whether the Buffa hypoxia signature or direct imaging modalities such as FMISO-PET can serve as clinical proxies for this threshold is an open question with direct translational relevance [32, 38].

The finding that hypofractionation achieved higher TCP than daily SCRT under equal total-dose conditions is an unexpected result that merits prospective experimental validation. The model predicts that this advantage is driven by repopulation suppression rather than per-cell radiosensitivity, a mechanistic hypothesis testable through in vitro clonogenic survival assays with timed re-irradiation at varying inter-fraction intervals.

## 5. Conclusion

This study demonstrates that spatial geometry of the tumour oxygen microenvironment is a determinant of immune clearance success following fractionated short-course radiotherapy in colorectal cancer. Using a spatially explicit hybrid agent-based model in which all cellular behaviours are governed by local oxygen tension and ATP availability, we showed that treatment outcome across independent realisations was predicted with complete separation by the oxygen context of surviving cells at end of treatment—not by surviving cell count alone. Cells persisting in the hypoxic core were simultaneously protected from radiation by reduced oxygen enhancement ratio and from immune killing by hypoxia-dependent CTL suppression, constituting a spatial refuge that the model identifies as the functional origin of treatment escape. Fractionation schedule analysis demonstrated that inter-fraction interval critically modulates repopulation dynamics and tumour control probability under equal total-dose conditions. Transcriptomic analysis of 284 Stage II/III CRC patients confirmed that significant intratumoral hypoxia is a clinically prevalent feature of the target disease, grounding the model’s oxygen assumptions in real patient biology. These findings suggest that the spatial distribution of hypoxia within the tumour microenvironment—quantifiable through imaging or transcriptomic surrogates—may represent a clinically actionable stratification criterion for predicting radiotherapy response and optimising treatment scheduling in locally advanced colorectal cancer.

## Data Availability

Simulation code and datasets are available at: https://doi.org/10.5281/zenodo.18646212

## Ethics Statement

This study did not involve human participants, human tissue, or animal experiments. Ethics approval and informed consent were not required. TCGA RNA sequencing data used in the transcriptomic analysis are publicly available through the Genomic Data Commons portal at portal.gdc.cancer.gov under open-access data policies. No additional patient-level data were generated in this study.

## Author Contributions

Sa Dharmasastha Karthikeya: Conceptualisation, Methodology, Software, Formal analysis, Investigation, Writing – Original Draft. Prathiba Jonnala: Writing – Review & Editing, Supervision.

## Declaration of Generative AI and AI-Assisted Technologies in the Writing Process

During the preparation of this manuscript, the authors used Claude (Anthropic) to support content organisation, language refinement, and structural editing of the manuscript text into L^A^T_E_X. The authors have reviewed and edited all AI-assisted content and take full responsibility for the accuracy, integrity, and originality of the work. All model development, simulation code, data analysis, figure generation, and scientific interpretation were performed entirely by the authors without AI assistance. AI tools were not used to generate or alter any figures, images, or data presented in this manuscript.

## References

[1] H. Sung, J. Ferlay, R. L. Siegel, M. Laversanne, I. Soerjomataram, A. Jemal, F. Bray, Global cancer statistics 2020: GLOBOCAN estimates of incidence and mortality worldwide for 36 cancers in 185 countries, CA: A Cancer Journal for Clinicians 71 (3) (2021) 209–249. doi:10.3322/caac.21660.

[2] A. Benson, A. Venook, M. M. Al-Hawary, M. A. Arain, Y.-J. Chen, K. K. Ciombor, et al., NCCN guidelines for colon cancer, Journal of the National Comprehensive Cancer Network 18 (3) (2020) 329–338. doi:10.6004/jnccn.2020.0014.

[3] K. Bujko, M. P. Nowacki, A. Nasierowska-Guttmejer, W. Michalski, M. Bebenek, M. Kryj, Long-term results of a randomized trial comparing preoperative short-course radiotherapy with preoperative conventionally fractionated chemoradiation for rectal cancer, British Journal of Surgery 93 (10) (2006) 1215–1223. doi:10.1002/bjs.5506.

[4] R. R. Bahadoer, E. A. Dijkstra, B. van Etten, C. A. M. Marijnen, H. Putter, E. M.-K. Kranenbarg, et al., Short-course radiotherapy followed by chemotherapy before total mesorectal excision (TME) versus preoperative chemoradiotherapy, TME, and optional adjuvant chemotherapy in locally advanced rectal cancer (RAPIDO): a randomised, open-label, phase 3 trial, The Lancet Oncology 22 (1) (2021) 29–42. doi:10.1016/S1470-2045(20)30555-6.

[5] F. Slevin, C. R. Hanna, A. Appelt, C. Cunningham, C. A. M. Marijnen, D. Sebag-Montefiore, R. Muirhead, The long and the short of it: the role of short-course radiotherapy in the neoadjuvant management of rectal cancer, Clinical Oncology 34 (5) (2022) e210–e217. doi:10.1016/j.clon.2022.01.014.

[6] S. Demaria, S. C. Formenti, Radiotherapy effects on anti-tumor immunity: implications for cancer treatment, Frontiers in Oncology 3 (2013) 128. doi:10.3389/fonc.2013.00128.

[7] A. D. Garg, D. Nowis, J. Golab, P. Vandenabeele, D. V. Krysko, P. Agostinis, Immunogenic cell death, DAMPs and anticancer therapeutics: an emerging amalgamation, Biochimica et Biophysica Acta (BBA) – Reviews on Cancer 1805 (1) (2010) 53–71. doi:10.1016/j.bbcan.2009.08.003.

[8] Z. Jin, F. A. Sinicrope, Mismatch repair-deficient colorectal cancer: building on checkpoint blockade, Journal of Clinical Oncology 40 (24) (2022) 2735–2750. doi:10.1200/JCO.21.02691.

[9] M. J. Overman, R. McDermott, J. L. Leach, S. Lonardi, H.-J. Lenz, M. A. Morse, et al., Nivolumab in patients with metastatic DNA mismatch repair-deficient or microsatellite instability-high colorectal cancer (CheckMate 142): an open-label, multicentre, phase 2 study, The Lancet Oncology 18 (9) (2017) 1182–1191. doi:10.1016/S1470-2045(17)30422-9.

[10] P. Vaupel, A. Mayer, M. Höckel, Impact of hemoglobin levels on tumor oxygenation: the higher, the better?, Strahlentherapie und Onkologie 182 (2) (2006) 63–71. doi:10.1007/s00066-006-1543-7.

[11] R. Sullivan, C. H. Graham, Hypoxia-driven selection of the metastatic phenotype, Cancer and Metastasis Reviews 26 (2) (2007) 319–331. doi:10.1007/s10555-007-9062-2.

[12] M. W. Dewhirst, Y. Cao, B. Moeller, Cycling hypoxia and free radicals regulate angiogenesis and radiotherapy response, Nature Reviews Cancer 8 (6) (2008) 425–437. doi:10.1038/nrc2397.

[13] M. C. Joiner, A. J. van der Kogel (Eds.), Basic Clinical Radiobiology, 4th Edition, CRC Press, 2009. doi:10.1201/b15450.

[14] B. D. Michael, A multiple-radical model for radiation action on DNA and the dependence of OER on LET, International Journal of Radiation Biology 69 (3) (1996) 351–358. doi:10.1080/095530096145799.

[15] P. Fan, N. Zhang, E. Candi, M. Agostini, M. Piacentini, F. Bernassola, et al., Alleviating hypoxia to improve cancer immunotherapy, Oncogene 42 (49) (2023) 3591–3604. doi:10.1038/s41388-023-02869-2.

[16] A. L. Doedens, A. T. Phan, M. H. Stradner, J. K. Fujimoto, J. V. Nguyen, E. Yang, et al., Hypoxia-inducible factors enhance the effector responses of CD8^+^ T cells to persistent antigen, Nature Immunology 14 (11) (2013) 1173–1182. doi:10.1038/ni.2714.

[17] R. A. Gatenby, R. J. Gillies, Why do cancers have high aerobic glycolysis?, Nature Reviews Cancer 4 (11) (2004) 891–899. doi:10.1038/nrc1478.

[18] M. G. Vander Heiden, L. C. Cantley, C. B. Thompson, Understanding the Warburg effect: the metabolic requirements of cell proliferation, Science 324 (5930) (2009) 1029–1033. doi:10.1126/science.1160809.

[19] M. V. Liberti, J. W. Locasale, The Warburg effect: how does it benefit cancer cells?, Trends in Biochemical Sciences 41 (3) (2016) 211–218. doi:10.1016/j.tibs.2015.12.001.

[20] S. D. Karthikeya, P. Jonnala, Spatiotemporal simulation of radiotherapy impact on Stage III sigmoid colon cancer using a metabolically coupled agent-based model, bioRxiv Preprint (2025). doi:10.1101/2025.04.19.649627.

[21] V. A. Kuznetsov, I. A. Makalkin, M. A. Taylor, A. S. Perelson, Nonlinear dynamics of immunogenic tumors: parameter estimation and global bifurcation analysis, Bulletin of Mathematical Biology 56 (2) (1994) 295–321. doi:10.1016/S0092-8240(05)80260-5.

[22] J. S. Lowengrub, H. B. Frieboes, F. Jin, Y.-L. Chuang, X. Li, P. Macklin, S. M. Wise, V. Cristini, Nonlinear modelling of cancer: bridging the gap between cells and tumours, Nonlinearity 23 (1) (2010) R1–R91. doi:10.1088/0951-7715/23/1/R01.

[23] J. J. Casciari, S. V. Sotirchos, R. M. Sutherland, Mathematical modelling of microenvironment and growth in EMT6/Ro multicellular tumour spheroids, Cell Proliferation 25 (1) (1992) 1–22. doi:10.1111/j.1365-2184.1992.tb01433.x.

[24] P. B. Vermeulen, D. Verhoeven, H. Fierens, G. Hubens, G. Goovaerts, E. Van Marck, E. A. De Bruijn, A. T. Van Oosterom, L. Y. Dirix, Microvessel quantification in primary colorectal carcinoma: an immunohistochemical study, British Journal of Cancer 71 (2) (1995) 340–343. doi:10.1038/bjc.1995.68.

[25] R. A. Cairns, I. S. Harris, T. W. Mak, Regulation of cancer cell metabolism, Nature Reviews Cancer 11 (2) (2011) 85–95. doi:10.1038/nrc2981.

[26] J. F. Fowler, The linear-quadratic formula and progress in fractionated radiotherapy, The British Journal of Radiology 62 (740) (1989) 679–694. doi:10.1259/0007-1285-62-740-679.

[27] D. J. Brenner, E. J. Hall, Fractionation and protraction for radiotherapy of prostate carcinoma, International Journal of Radiation Oncology, Biology, Physics 43 (5) (1999) 1095–1101. doi:10.1016/S0360-3016(98)00438-6.

[28] S. M. Bentzen, M. C. Joiner, The linear-quadratic approach in clinical practice, in: Basic Clinical Radiobiology, 5th Edition, CRC Press, 2018, pp. 112–124. doi:10.1201/9780429490606.

[29] J. Jiménez-Sánchez, Á. Martínez-Rubio, A. Popov, J. Pérez-Beteta, Y. Azimzade, D. Molina-García, et al., A mesoscopic simulator to uncover heterogeneity and evolutionary dynamics in tumors, PLOS Computational Biology 17 (2) (2021) e1008266. doi:10.1371/journal.pcbi.1008266.

[30] J. H. Goldie, A. J. Coldman, A mathematic model for relating the drug sensitivity of tumors to their spontaneous mutation rate, Cancer Treatment Reports 63 (11–12) (1979) 1727–1733.

[31] A. Saltelli, P. Annoni, I. Azzini, F. Campolongo, M. Ratto, S. Tarantola, Variance based sensitivity analysis of model output: design and estimator for the total sensitivity index, Computer Physics Communications 181 (2) (2010) 259–270. doi:10.1016/j.cpc.2009.09.018.

[32] F. M. Buffa, A. L. Harris, C. M. West, C. J. Miller, Large meta-analysis of multiple cancers reveals a common, compact and highly prognostic hypoxia metagene, British Journal of Cancer 102 (2) (2010) 428–435. doi:10.1038/sj.bjc.6605450.

[33] M. A. Swartz, M. E. Fleury, Interstitial flow and its effects in soft tissues, Annu. Rev. Biomed. Eng. 9 (1) (2007) 229–256.

[34] H. R. Withers, J. M. G. Taylor, B. Maciejewski, The hazard of accelerated tumor clonogen repopulation during radiotherapy, Acta Oncologica 27 (2) (1988) 131–146. doi:10.3109/02841868809090333.

[35] C. Capirci, V. Valentini, L. Cionini, A. De Paoli, C. Rodel, R. Glynne-Jones, et al., Prognostic value of pathologic complete response after neoadjuvant therapy in locally advanced rectal cancer: long-term analysis of 566 ypcr patients, International Journal of Radiation Oncology, Biology, Physics 72 (1) (2008) 99–107. doi:10.1016/j.ijrobp.2007.12.022.

[36] H. Wang, H. Jiang, M. Van De Gucht, M. De Ridder, Hypoxic radioresistance: can ROS be the key to overcome it?, Cancers 11 (1) (2019) 112. doi:10.3390/cancers11010112.

[37] P. Vaupel, A. Mayer, M. Höckel, Tumor hypoxia and malignant progression, in: Methods in Enzymology, Vol. 381, Academic Press, 2004, pp. 335–354. doi:10.1016/S0076-6879(04)81023-9.

[38] M. R. Horsman, J. Overgaard, The impact of hypoxia and its modification of the outcome of radiotherapy, Journal of Radiation Research 57 (S1) (2016) i90–i98. doi:10.1093/jrr/rrw007.

[39] G. G. Steel, T. J. McMillan, J. H. Peacock, The 5Rs of radiobiology, International Journal of Radiation Biology 56 (6) (1989) 1045–1052. doi:10.1080/09553008914552491.

[40] A. B. Sharabi, P. T. Tran, M. Lim, C. G. Drake, T. L. DeWeese, Stereo-tactic radiotherapy combined with immunotherapy: augmenting radiation’s role in local and systemic treatment, Oncology 29 (5) (2015) 331–340.

[41] S. Jafari Nivlouei, M. Soltani, J. Carvalho, R. Travasso, M. R. Salimpour, E. Shirani, Multiscale modeling of tumor growth and angiogenesis: evaluation of tumor-targeted therapy, PLOS Computational Biology 17 (6) (2021) e1009081. doi:10.1371/journal.pcbi.1009081.

[42] G. G. Powathil, M. A. J. Chaplain, A hybrid multiscale approach in cancer modelling and treatment prediction, in: Mathematical Oncology 2013, Springer New York, 2014, pp. 237–263. doi:10.1007/978-1-4939-0458-79.

